# A manifold neural population code for space in hippocampal coactivity dynamics independent of place fields

**DOI:** 10.1101/2021.07.26.453856

**Authors:** Eliott R.J. Levy, Simón Carrillo-Segura, Eun Hye Park, William T. Redman, José R. Hurtado, SueYeon Chung, André A. Fenton

**Affiliations:** Center for Neural Science, New York University, New York, NY 10003, USA; Graduate Program in Mechanical and Aerospace Engineering, Tandon School of Engineering, New York University, Brooklyn, NY 11201; Interdepartmental Graduate Program in Dynamical Neuroscience, University of California, Santa Barbara, Santa Barbara, CA 93106, USA; Flatiron Institute Center for Computational Neuroscience, New York, NY 10010. USA; Neuroscience Institute at the NYU Langone Medical Center, New York, NY 10016, USA

**Author notes:** Correspondence: André Fenton, Neurobiology of Cognition Laboratory, Center for Neural Science, New York University, 4 Washington Place, Room 980. Equal Contribution (Eliott R.J. Levy, Simón Carrillo-Segura).

**Keywords:** Keywords: Hippocampus, place cell, cell assembly, neural synchrony, synapsemble, environment, context, remapping, reregistering

## Abstract

Hippocampus is comprised of ∼20% place cells, discharging in cell-specific locations (“place fields”), standardly interpreted as a dedicated neuronal code for space. However, place cell discharge is temporally unreliable across seconds and days, and place fields are multimodal, suggesting an alternative “ensemble cofiring” spatial code with manifold dynamics that does not require reliable spatial tuning. We evaluated these hypotheses using GCaMP6f and miniature microscopes to image mouse CA1 ensemble activity in two environments, across 3 weeks. Both place fields and ensemble coactivity relationships appear to “remap,” being distinct between, and (weakly) similar within environments. Decoding location as well as environment from 1-s ensemble location-specific discharge is effective and improves with experience. Decoding the environment (but not location) from cell-pair coactivity relationships is also effective and improves with experience, even after removing place tuning. Discriminating environments from 1-s ensemble coactivity relies crucially on the cells with the most anti-cofiring cell-pair relationships because ensemble activity is internally-organized on a low-dimensional manifold of non-linear cofiring relationships that intermittently reregisters to environments according to the anti-cofiring subpopulation activity.

## Introduction

The hippocampus is crucial for spatial navigation as well as memory, and although hippocampal place cells are tuned to discharge in cell-specific locations within an environment called “place fields,” their relationship to spatial cognition is unclear (Muller et al., 1987; O’Keefe, 1976, 1979). When animals change environments, the location-specific tuning of each place cell changes uniquely, a phenomenon called “remapping” (Fig. 1A). Remapping means that the ensemble pattern of cofiring is unique and environment-specific, constituting a spatial code that relates neural activity to the environment (Alme et al., 2014; Bostock et al., 1991; Colgin et al., 2008; Kubie et al., 2020; Muller and Kubie, 1987; Wills et al., 2005).

**Figure 1.**
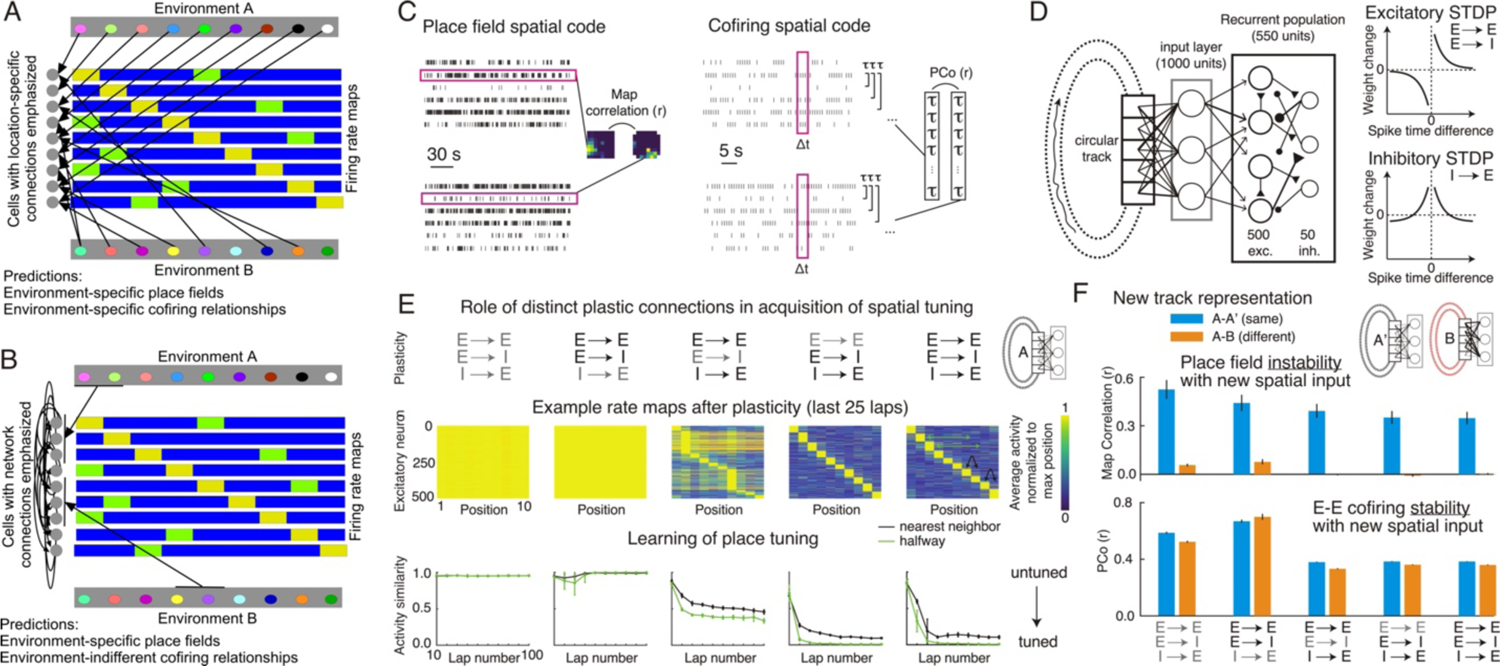
Neural coding hypotheses for representing environments. (A) Schematic illustrating the standard concept of remapping that emphasizes that cell-specific location-specific inputs change both place field locations and ensemble cofiring relationships across environments, and (B) the concept of reregistration that emphasizes intrinsic hippocampal inputs that do not change between environments despite a change in the registration of environmental information onto the mostly invariant network cofiring relationships. The firing rate maps illustrate environment-specific multipeaked place fields (yellow and green). (C) Illustration of hypothesized (left) dedicated place field and (right) ensemble cofiring codes for environments, and how they can be quantified, by single cell spatial firing stabilities, or by computing the stability (PCo) of the vector of cell pair correlations, respectively. PCo computes a Pearson correlation (r) to measure the similarity between the set of n(n-1)/2 Kendall correlations (τ) that describe the pairwise activity correlations within an n-cell ensemble (Neymotin et al., 2017). (D) Schematic of neural network encoding of a circular environment with STDP learning rules. (E, top) Learned network representation of the circular environment after enabling (black) or disabling (grey) STDP learning at cell-type specific connections, as measured on (middle) time-averaged rate maps and by (bottom) spatial activity similarity. The nearest neighbor metric measures the difference between the peak activity and the activity at the closest positions to the location of the peak; the halfway metric measures the difference between the peak activity and the activity at the position halfway across the track. (F) Consequence of remapping the external inputs to the network (track B) compared to an unchanged mapping (null remapping, track A’). Evaluation of (top) dedicated place field and (bottom) ensemble cofiring codes in the track A and B environments as a function of enabled (black) or disabled (grey) STDP learning at cell-type specific connections. Discrimination between environments is robust with the place field code (top) and is most sensitive to inhibitory plasticity. This is especially prominent in the ensemble cofiring code (bottom) that in contrast to place firing, is relatively similar across experiences of the same and different environments.

Although the concept of remapping means a rearrangement of temporal cofiring relationships, the vast majority of studies have inferred that a rearrangement of cofiring relationships has occurred by only measuring that the spatial tuning, specifically the time-normalized place fields of individual cells has changed. It is, however, possible that the place fields of individual cells change without substantially changing their short-timescale cofiring relationships. It is especially easy (but not necessary) to imagine how this can happen if one accepts that most place cells have multiple non-periodic place fields, as is observed in environments with linear dimensions greater than a meter (Fenton et al., 2008; Harland et al., 2021). As Fig 1A illustrates, remapping implies that place fields are determined by environmental features like geometry and orienting cues (Hetherington and Shapiro, 1997; O’Keefe and Burgess, 1996; Olypher et al., 2003). In contrast, it is possible to observe changed place fields, the standard evidence of remapping, when only the registration changes between the environment and a population of invariant cofiring relationships that are internally-organized (Fig. 1B). Whether remapping is actually reregistration has important implications not only for understanding the hippocampal space code and navigation, but also for understanding memory.

Because both memory and environmental context representations rely on the hippocampus, the dominant hypothesis asserts that remapping and memory are intimately linked; changes in place tuning are assumed to correspond to changes in memory representations (Guzowski et al., 2004; Leutgeb et al., 2005b; Leutgeb et al., 2005c; Lever et al., 2002; Wills et al., 2005), in particular, episodic memory representations that include information about environments (Kentros, 2006; Mizumori, 2006). We note however, that it is not straightforward how the arrangement of place fields can represent a particular environment at the millisecond-to-second timescale of neural computations without initial, extensive spatial exploration (McHugh and Tonegawa, 2007), and for a number of additional fundamental reasons: 1) place cells are multimodal, having multiple place fields in environments bigger than about 1m^2^ (Fenton et al., 2008; Harland et al., 2021); 2) discharge in firing fields is extremely variable during the 1-to-5 seconds it takes to cross the place field (Fenton et al., 2010; Fenton and Muller, 1998; Jackson and Redish, 2007); 3) the firing rate in a place field varies systematically across behavioral episodes, called “rate remapping” (Leutgeb et al., 2005b); and 4) only a minority of place cells have stable place fields across days in familiar environments (Lee et al., 2020; Ziv et al., 2013). The consequence of cells with multiple place fields demonstrates that this alone can degrade the ability to decode environments from ensemble discharge (Fig. S1). We also note that studies that tested the hypothesis that remapping is related to memory have been remarkably unsupportive (Duvelle et al., 2019; Jeffery et al., 2003; Leutgeb et al., 2005a; Moita et al., 2004; van Dijk and Fenton, 2018).

One possible reason for the inability to relate remapping to memory could be incorrect assumptions about how neural information is encoded; indeed, what is the nature of the neural code? Perhaps the code for environments is different from the code for locations (Tanaka et al., 2018). Studies standardly average the activity of individual cells over minutes, discarding discharge fluctuations in time as well as any cofiring relationships with other cells to extract the cell’s relationship to a certain variable, such as its location-specific tuning. Such analyses are performed under the (incorrect) assumption that place-discharge relationships are in a steady state, although they are not (Dvorak et al., 2018; Fenton et al., 2010; Fenton and Muller, 1998; Jackson and Redish, 2007; Kay et al., 2020). This standard approach, a dedicated-rate place field hypothesis assumes the cell’s momentary firing rate independently carries information that can be adequately extracted by analysis of each cell’s place tuning. The firing relations of each cell to other neurons are assumed to be uninformative, or at least secondary, which is why they are explicitly ignored by data representations like the firing rate maps that define place fields. Indeed, it is intuitive to imagine that place cells with overlapping fields might be expected to cofire because time and space are confounded. However, whereas some place cell pairs with overlapping place fields reliably cofire, other pairs do not, and yet other pairs discharge independently on the ms-to-s timescales of crossing firing fields and neural computation (Fig. S3; Fenton, 2015a; Harris et al., 2003; Kelemen and Fenton, 2013). This motivates an alternative cofiring (ensemble or cell assembly) coding hypothesis that asserts information is encoded in the momentary cofiring patterns of activity that are expressed by large groups of neurons -- each cofiring pattern encodes different information (Harris et al., 2003; Meshulam et al., 2017; Stefanini et al., 2020). Such cofiring neural codes are explicit in the increasingly popular recognition that high-dimensional neural population activity (each dimension being a neuron) can be constrained to a manifold, a low-dimensional sub-space defined by recurring across-cell patterns of coactivity (Babichev et al., 2016; Chaudhuri et al., 2019; Dabaghian et al., 2014; Ebitz and Hayden, 2021; Gardner et al., 2022; Jazayeri and Afraz, 2017; Low et al., 2022; McNaughton et al., 2006; Park et al., 2019; Peyrache et al., 2015; Rubin et al., 2019; Umakantha et al., 2021). Note that higher-order firing patterns constrained on a manifold carry information, and that in principle, without changing shape, such manifolds can describe infinitely many individual cell discharge patterns and even an infinity of lower-order cofiring patterns (Dabaghian et al., 2014; Heeger and Mackey, 2019; Jazayeri and Afraz, 2017), as recently demonstrated for MEC grid cells (Gardner et al., 2022; Low et al., 2022). Such discharge patterns are constrained by “on-manifold” cofiring relationships and because they need not ever repeat, they need not depend on tuning to external variables like locations. Here we repeated a standard place cell remapping procedure to evaluate the dedicated place field and ensemble cofiring hypotheses for representing distinct environments in the hippocampus.

## Results

### Distinguishing place tuning and cofiring contributions to representing environments: a neural network model

We formalized the place field and cofiring hypotheses by examining a heuristic neural network model designed to evaluate the contributions of place tuning (Fig. 1C, left) and cofiring (Fig. 1C, right) to representing environments. The recurrently connected excitatory-inhibitory network receives excitatory input from units tuned to randomly sampled positions of a circular track (Fig. 1D, left). The recurrent network weights change according to standard spike-timing dependent plasticity (STDP) rules (Fig. 1D, right) that are sufficient to generate place-selective activity (place fields, Fig. 1E, middle) in the recurrent network after ∼30 laps on the track (Fig. 1E, bottom). Among other features, simply enlarging the track generates multiple firing fields once there has been sufficient time for exploration and the learning rules to operate, as is observed in real environments (Fenton et al., 2008; Harland et al., 2021). Location-specificity could be assessed by the dissimilarity of the place fields (Fig. 1E), as well as the ability to decode current position from momentary network activity (data not shown). Remarkably, E→I and I→E plasticity are necessary for developing the place tuning, but E→E plasticity is not (Fig. 1E), reproducing observations after blocking NMDA receptor-dependent LTP (Kao et al., 2017; Kentros et al., 1998; Lamsa et al., 2005), which dominates at E→E connections but not at E→I or I→E connections in hippocampus (Lamsa et al., 2005). This illustrates that tunable excitation-inhibition coordination can be important for network and memory function, as observed (Caroni, 2015; Chung et al., 2021; Dvorak et al., 2021; Fenton, 2015b; Lamsa et al., 2010; Mongillo et al., 2018; van Dijk and Fenton, 2018); see papers dedicated to the topic (Kullmann and Lamsa, 2011). To simulate remapping after learning a second environment, we randomized the position-to-input relationships and allowed the recurrent weights to change according to the same STDP rules during another 30 laps. As expected, the place fields of the network units changed; they remapped the new track in the sense that the spatial firing of each cell is unrelated across the two tracks (Fig. 1F, top; Fig. S2A). In contrast, the population of pairwise E-E cofiring relationships behaves differently, demonstrating it is possible to change firing fields (the conventional measure of remapping) without substantially changing cofiring relationships (Fig. S2). Cofiring relationships are marginally more stable when spatial inputs are unchanged compared to when they are changed (i.e. remap). Cofiring relationships tend to change with experience, regardless of whether the environmental inputs change, and these changes are most strongly dependent on I→E plasticity and they are required for location-specific activity, in this model. This network simulation is a heuristic, demonstrating that several key ideas are feasible: 1) that a place field code and a cofiring code could be distinct and operate in parallel; 2) that place field remapping between environments may occur without changing the majority of E-E cofiring relationships in a network; 3) that remapping may be a misnomer for what is a reregistration between an internally-rigid neural activity representation and external inputs; 4) that place field remapping/reregistration depends on the environment-specific anti-cofiring E-E relationships; a consequence of necessarily altered I→E (and E→I) functional connections, which have been observed as experience-dependent (Mongillo et al., 2018) and learning-dependent changes in network inhibition (Chung et al., 2021). If correct, these ideas will require changes to standard ideas, analyses, and interpretations of the spatial tuning of hippocampal neurons, and perhaps cells in other brain regions too (Ebitz and Hayden, 2021).

### Calcium imaging in CA1

To examine the validity of dedicated place field and ensemble cofiring hypotheses, we infected mouse CA1 with a virus expressing GCaMP6f (Cai et al., 2016; Chen et al., 2013). While the mice were freely exploring, we recorded CA1 ensemble activity with a miniature microscope (http://miniscope.org; Cai et al., 2016) placed over a chronically implanted gradient-index (GRIN) lens (Ziv et al., 2013). During the recordings, the animals (n=6) explored two physically distinct environments, a cylinder and box. The mice were habituated to the miniscope in their home cage and only experienced the two environments during recordings according to the protocol in Fig. 2A. Raw calcium traces were extracted and segmented from videos using the CNMF-E algorithm (Pnevmatikakis et al., 2016; Zhou et al., 2018), and, when possible, registered to individual cells across the three-week protocol (Fig. 2B). Over 50% of individual cells could be identified two weeks apart, though not unsurprisingly the registration was greater across shorter intervals (Fig. 2C). Raw calcium traces were converted with the CNMF-E algorithm to spiking activity for analysis (Fig. 2D).

**Figure 2.**
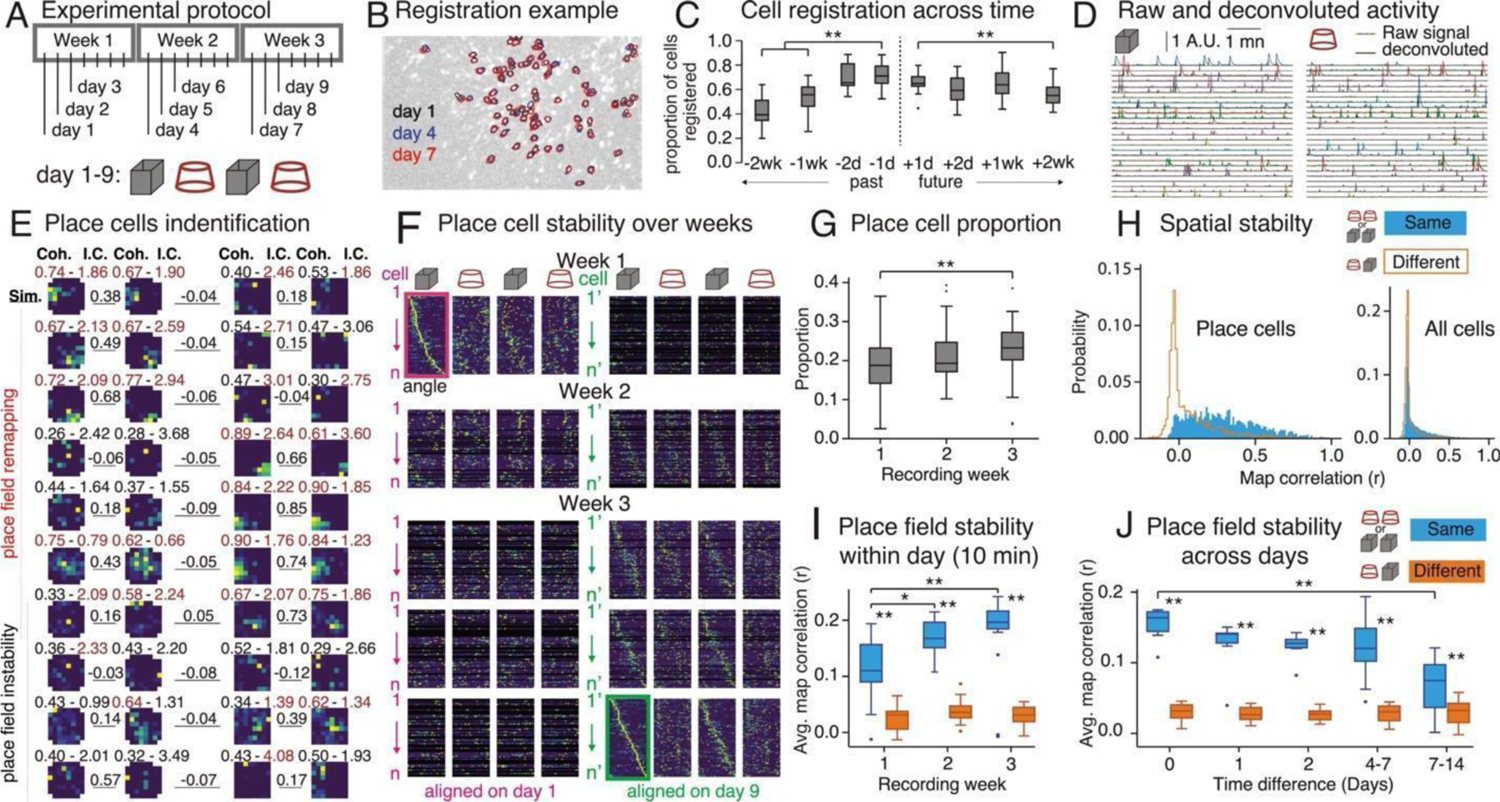
Location-specific CA1 activity imaged across weeks is transient, describes a minority of the CA1 population, and remaps between environments. (A) Experimental protocol. (B) Example and (C) quantification of cell registration across weeks (Past: Time F_3,187_ = 56.89, p= 10^-26^,η^2^ = 0.48; Dunnett’s tests: −1 day > −2, −1 week(s); Future: Time F_3,187_ = 7.31, p= 10^-4^,η^2^ = 0.12; Dunnett’s tests: 1 day > 2 weeks). (D) Activity time series in two environments illustrated by a single 25-cell example from a 546-cell ensemble. € Activity rate map examples from 10 cells from a 588-cell ensemble across two environments on recording day 7. Place field quality metrics, spatial coherence (Coh.; left) and information content (I.C.; right) are indicated above each map, red indicating significance compared to shuffled; average rate map similarity (Sim.; r) is indicated between maps. (F) Linearized ensemble activity rate maps across three weeks (week 1: day 1; week 2: day 4; week 3: days 7,8,9), matched and aligned to activity on recording day 1 (left, 312 cells) and recording day 9 (right, 460 cells). (G) Proportions of cells with significant spatial coherence and information content, classified as place cells (Week: F_2,64_ = 43.70, p= 10^-12^,η^2^ = 0.085; Dunnett’s tests: week 1 < week 3). (H, left) Place map similarity across environments on weeks 2 and 3 illustrates remapping of place cells but (right) not the CA1 population. (I) Place map similarity within day improves across weeks (Week: F_2,28_ = 20.02, p = 10^-6^, η^2^ = 0.051; Env. Change: F_2,29_ = 80.55, p = 10^-10^, η^2^=0.65; Interaction: F_2,28_ = 4.83, p = 0.016, η^2^=0.026; Dunnett’s tests: week 1 < weeks 2, 3 for similar environments; post-hoc: Same vs. Different is significant each week, **p’s ≤ 10^-5^). (J) Place map similarity degrades across days (Time difference: F_4,132_ = 12.68, p = 10^-9^, η^2^=0.11; Env. Change: F_1,132_ = 228.37, p = 10^-30^, η^2^=0.53; Interaction: F_4,132_ = 9.32, p = 10^-6^,η^2^=0.12; Dunnett’s tests: day 0 > 7-14 days for similar environments; post-hoc: Same vs. Different is significant at each of the five-time intervals, **p’s ≤ 10^-3^).

### Place field measurements of remapping

In each environment, cells expressed place fields (Fig. 2E) that were stable within a day and unstable across days as previously reported (Lee et al., 2020; Ziv et al., 2013). To better visualize the organization of the ensemble place response, the environment was linearized by plotting activity rates as a function of angle (Fig. 2F). By inspection, the location-specific pattern expressed in week 3 (aligned on recording day 9) was observed during week 2, suggesting the location-specific activity was more stable than it was in week 1 (aligned on recording day 1). These observations suggest that place tuning was learned during the first week. While consistently constituting a minority of the population (Stefanini et al., 2020; Wilson and McNaughton, 1993), the percentage of place cells increased with experience from 20% to 25% (Fig. 2G). Each day, place cells expressed stable firing fields between trials in the same environment, but location-specificity changed between trials of different environments (Fig. 2H, left). Such remapping is not population wide as the measurable difference vanishes when all recorded cells are considered. This is largely because the place tuning of non-place cells is unstable between any pair of trials (Fig. 2H, right). The average within-day place field stability of place cells grows with experience from 0.1 during week 1 to 0.2 during week 3 (Fig. 2I), about 6 times larger than for distinct environments on week 3. Yet, this place field stability degrades across days (Fig. 2J), as previously described (Lee et al., 2020; Ziv et al., 2013). These measurements demonstrate standard mouse place cell phenomena in the data set.

### Ensemble cofiring measurements of remapping between environments

Next, we examined the change in CA1 activity across the two environments without regard to positional information. To evaluate whether distinct environments can be encoded by conjoint discharge, we measured the correlations of cell pairs, to estimate the recurrence of higher-order correlations, responsible for the ensemble’s characteristic population geometry (Fig. S3; Schneidman et al., 2006). In each trial, for each cell pair, we measured the coactivity of their 1-s activity time series by computing their Kendall correlation (Fig. 3A). To evaluate their stability across environments, we measured the population coordination (PCo) as the similarity of all tau values, computed as the Pearson correlation between two vectors of pairwise tau values (Fig. 3B; Neymotin et al., 2017; Talbot et al., 2018). PCo between trials of the same environments is three times higher than between distinct environments (Fig. 3C), indicating that ensemble coactivity discriminates environments. The individual cell pairs that differ the most between environments are the positively (τ > 0.3) or negatively (τ < −0.05) coactive cell pairs (Fig. 3A). PCo increases with experience (Fig. 3C, left) and deteriorates with greater time between the recordings (Fig. 3C, right), like place field measurements of remapping (Fig. 2I, J). These findings indicate that environments are discriminatively encoded in the 1-s coactivity relationships of CA1 neural activity.

**Figure 3.**
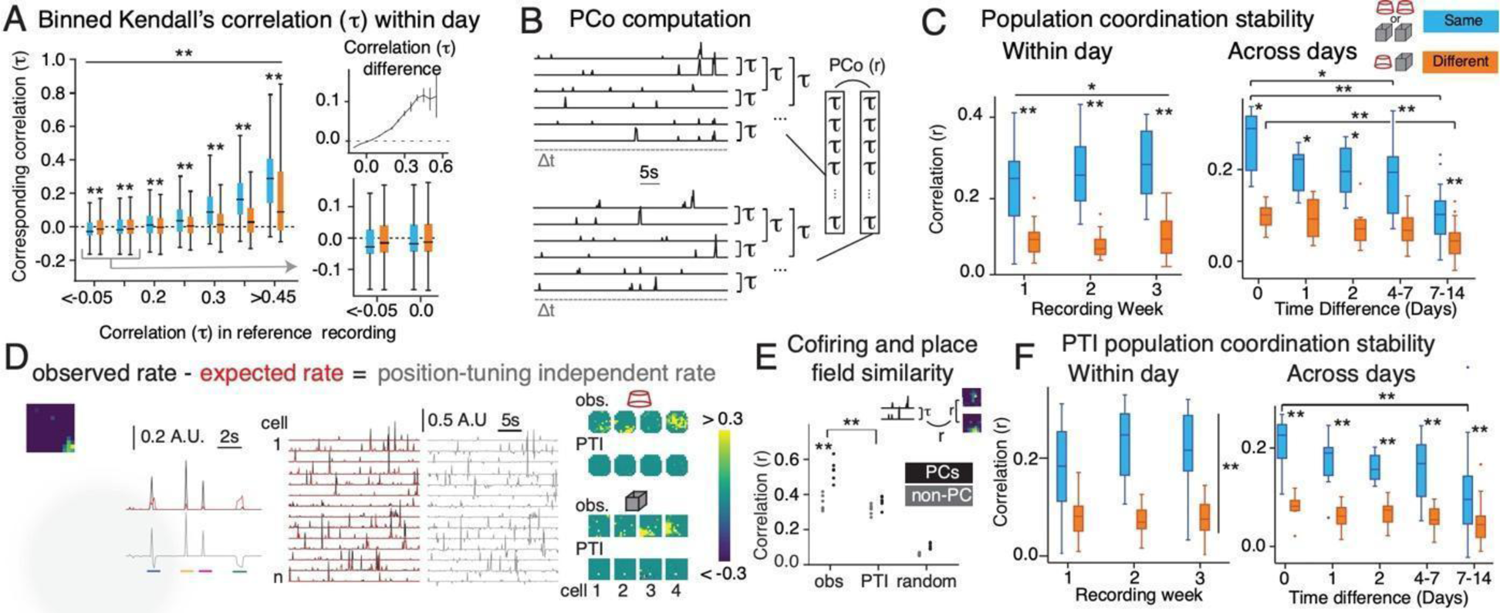
Ensemble cofiring distinguishes two environments independently of place tuning. (A) Comparison of coactivity (τ) stability of all cell pairs, from weeks 2 and 3, across same-day recordings, for different initial coactivity values. This illustrates that coactive cell pairs tend to remain coactive across the same, but not different environments (Coactivity Level: F_6,3285501_ = 9825.46, p = 0.00, η^2^ = 0.018; Env. Change: F_1,3285501_ = 9.82, p = 0.0017, η^2^ = 10^-6^; Interaction: F_6,3285501_ = 3395.58, p = 0.00, η^2^=0.0061; post-hoc: Same vs. Different is significant at each coactivity level, **p’s < 10^-15^) and (bottom inset) anti-coactive cell pairs remain anti-coactive. (top inset) Average signed differences confirm the effect of environment change. (B) Illustration of Population Coordination (PCo) calculation from cofiring distributions. The time intervals (Δτ = 1 s) used to discretize activity are illustrated as grey dashes, to scale. (C, left) Within-day coactivity stability improves across weeks (Week: F_2,27_ = 4.23, p = 0.025, η^2^=0.015; Env. Change: F_1,28_ = 92.32, p = 10^-10^, η^2^=0.56; Interaction: F_2,27_ = 0.99, p = 0.38, η^2^=0.022; post-hoc: Same vs. Different is significant each week, **p’s < 10^-4^). (C, right) Across-day cofiring stability degrades over time (Time Difference: F_4,132_ = 16.79, p = 10^-11^, η^2^=0.21; Env. Change: F_1,132_ = 94.66, p = 10^-17^, η^2^=0.32; Interaction: F_4,132_ = 4.04, p = 0.004, η^2^=0.072; Dunnett’s tests vs day 0, *p<0.0058, **p ≤ 10^-5^; post-hoc: Same vs. Different is significant at each time interval, **p ≤ 10^-3^). (D) Illustration that subtracting expected position-tuning from observed activity captures activity fluctuations independent of place tuning. (Left to right) Example activity map in the square environment and magnified version with four passes across the place field (star = path start). Observed, expected and PTI activity times series with the color-coded passes marked and additional examples from 14 cells. Observed and PTI activity maps from 4 Fig. 1 example cells. (E) Correlation between place field similarity and coactivity, computed from observed, PTI and randomized PTI activities, for place cells and non-place cells (Time-Series Type: F_2,30_ = 315.13, p = 10^-21^, η^2^ = 0.84; Functional Cell Class: F_1,30_ = 56.74, p = 10^-8^, η^2^ = 0.076; Interaction: F_2,30_ = 14.72, p = 10^-5^, η^2^ = 0.039; post-hoc: PC vs. non-PC for obs and obs vs. РТI for Place Cells significant, **p ≤ 10^-4^). (F, left) Within-day PTI coactivity stability across weeks (Week: F_2,27_ = 2.28, p = 0.12, η^2^ = 0.009; Env. Change: F_1,28_ = 46.67, p = 10^-7^, η^2^ = 0.52; Interaction: F_2,27_ = 2.38, p = 0.11, η^2^ = 0.028; post-hoc: Same vs. Different: t_67.44_ = 10.14, p = 10^-15^). (F, right) Across-day PTI coactivity stability also degrades over time (Time Difference: F_4,132_ = 5.23, p = 10^-4^, η^2^ = 0.088; Env. Change F_1,132_ = 66.57, p = 10^-13^, η^2^ = 0.33; Interaction: F_4,132_ = 1.69, p = 0.16, η^2^ = 0.079; Dunnett’s tests: day 0 > days 7-14 for similar environments, **p = 0.0050; post-hoc Same vs. Different significant at each time interval, **p’s < 0.008).

### Ensemble cofiring measurements of remapping between environments independently of position tuning

Because CA1 activity is influenced by the animal’s position, we evaluated the extent to which position tuning and overlapping place fields explains the 1-s coactivity relationships of CA1 activity. We calculated a position-tuning independent (PTI) rate by subtracting the expected rate from the cell’s observed rate, as schematized in Fig. 3D. The expected rate was computed from the position of the animal and the session-averaged firing-rate map (Fenton et al., 2010; Fenton and Muller, 1998). As a direct consequence, place fields computed from PTI rate disappear (Fig. 3D, right). Nonetheless, correspondence is high (0.4 < r < 0.6) between cell-pair correlations computed from the observed rates of the cell pairs and those computed from the PTI rates of these pairs, indicating activity correlations in cofiring that exist beyond position-tuning (Fig. S4B), as also observed ubiquitously in neocortex with cells tuned to different variables (Churchland et al., 2010; Hennequin et al., 2018; Lee et al., 1998; Schneidman et al., 2003). We stress that our use of PTI is not to claim that position tuning has no meaning (it signals position), but rather to assess what information is in cofiring independent of position tuning (see Fig. S6).

The cell-pair correlations computed from observed rates covary with spatial firing similarity for both place cells and non-place cells. The cell-pair correlations computed from PTI also covary with spatial firing similarity for both place cells and non-place cells, but at the same level as calculated with the observed rates for non-place cells (Fig. 3E). Taken together, this indicates that overlapping place fields are responsible for a small fraction of the relationship between cell-pair correlation and spatial firing similarity in place cells (Fig. 3E). PCo computed from PTI (PTI-PCo) decreases by approximately 20%, remains higher for trials between the same environment compared to between different environments, but does not change from week 1 to week 3 (Fig. 3F, left). Like PCo computed from observed rates (Fig. 3C, right), PTI-PCo also degrades over time (Fig. 3F, right). These findings indicate that the information contained in the cell’s position tuning is, in theory, not necessary for discriminating two environments using the activity of CA1 principal cells, including place cells. This motivated us to examine how coactivity relationships could be used for representing spatial information.

PTI coactivity; (top) PTI coactivity, computed every 60 s, is projected with SVM weights onto a single dimension that separates the two environments. (Bottom) Cell-pair coactivity from an example ensemble ordered by SVM weight each week; weights’ absolute value, indicative of separation strength, are classified by decile. (B) Cross-validated decoding accuracy is very high for all ensembles. (C) Difference between overall and first decile distribution of cell pair correlations (example distributions in inset and shown in purple in main figure) indicates importance of coactive and anti-coactive cell pairs, confirmed by (D) 2-D distribution of SVM weights against PTI coactivity. (E) Proportion of individual cells present in at least one cell pair in each SVM-ordered cell pair decile. (F) Difference between random and first decile, normalized distributions of individual cell participation (example distributions in inset and shown in purple in main figure) indicates overrepresentation of some cell in the first decile. (G) PTI network consistency normalized by the standard deviation of the cell-pair randomized distribution (Week: **F_2_,_6ū_** = 15.76, p=10^-6^, η^2^ = 0.094; Dunnett’s tests: week 0 < weeks 2, 3).

### Decoding environments from position-tuning independent ensemble coactivity

We then investigated whether the current environment could be reliably decoded from the PTI ensemble coactivity, from which we eliminated all position tuning, in order to discover whether there is a crucial contribution of coactivity to decoding. Coactivity was computed at 1-s resolution over the period of a minute, yielding up to five independent estimates of coactivity per trial. We trained a support-vector machine (SVM) decoder to project the coactivity along a single composite dimension that separates the two environments (Fig. 4A). The decoder almost always correctly identifies the current environment (Fig. 4B), demonstrating that the coactivity itself carries discriminative information, independent of position tuning. Remarkably, we find that the most discriminative cell pairs are either strongly coactive or strongly anti-coactive (Fig. 4C, D). Most individual cells contributed to the cell pairs in each decile (Fig. 4E), but some cells are overrepresented in the subset of discriminative cell pairs that constitutes the first decile (Fig. 4F). Taken together, these two characteristics of cell-pair activity co-fluctuations beyond positional tuning indicate that population correlation dynamics can be described as scale-free, strikingly similar to models that describe flocks of birds (Hemelrijk and Hildenbrandt, 2015). Indeed, power laws fit the distribution of activity correlations as a function of distance between the imaged soma; the best fit exponents are negative and slightly smaller than 1 (average ± s.d.: −0.90 ± 0.05; see Fig. S4D). Accordingly, the correlation length (distance at which the average correlation = 0) is large, indicating that on average, the activity of each cell non-causally influences the activity of every other cell. The environment-discriminating activity of the neural population is driven by a subset of the individual cells but impacts the whole population, which is reminiscent of the flocking and schooling behaviors of birds and fish (Hemelrijk and Hildenbrandt, 2011, 2012; Reynolds, 1987). We consequently calculated the network consistency (van Dijk and Fenton, 2018) as an estimate of the overall correspondence between cell pairs’ *(i* and *j)* PTI coactivity (τ_i,j_) and the pairs’ instantaneous PTI rate (η and η). Network consistency increases with experience (Fig. 4G) indicating an increased alignment of the 1-s short- and 300-s long-timescales of cell pair cofiring in the network. Repeating the set of coactivity calculations without first eliminating the place fields yielded similar, but less discriminative results (Fig. S4, Week: F_2,19_ = 34.96, p = 10^-7^, η^2^ = 0.023; Time-Series Type: F_1,20_ = 14.66, p = 0.001, η^2^ = 0.35; Interaction: F_2,19_ = 26.61, p = 10^-6^, η^2^ = 0.011; post-hoc: PTI > observed on weeks 1 and 3, p ≤ 0.004; p = 0.027 on week 2). Тhis is consistent with the possibility that the environment-specific information signaled by the coactivity of cells carries a separate type of information than the positional tuning measured by firing fields (Huxter et al., 2003). Parenthetically, the ability to decode current location from 1-s place cell activity vectors is somewhat better than the ability to decode current location from 1-s cofiring assessed by coincidence (Fig. S6).

**Figure 4.**
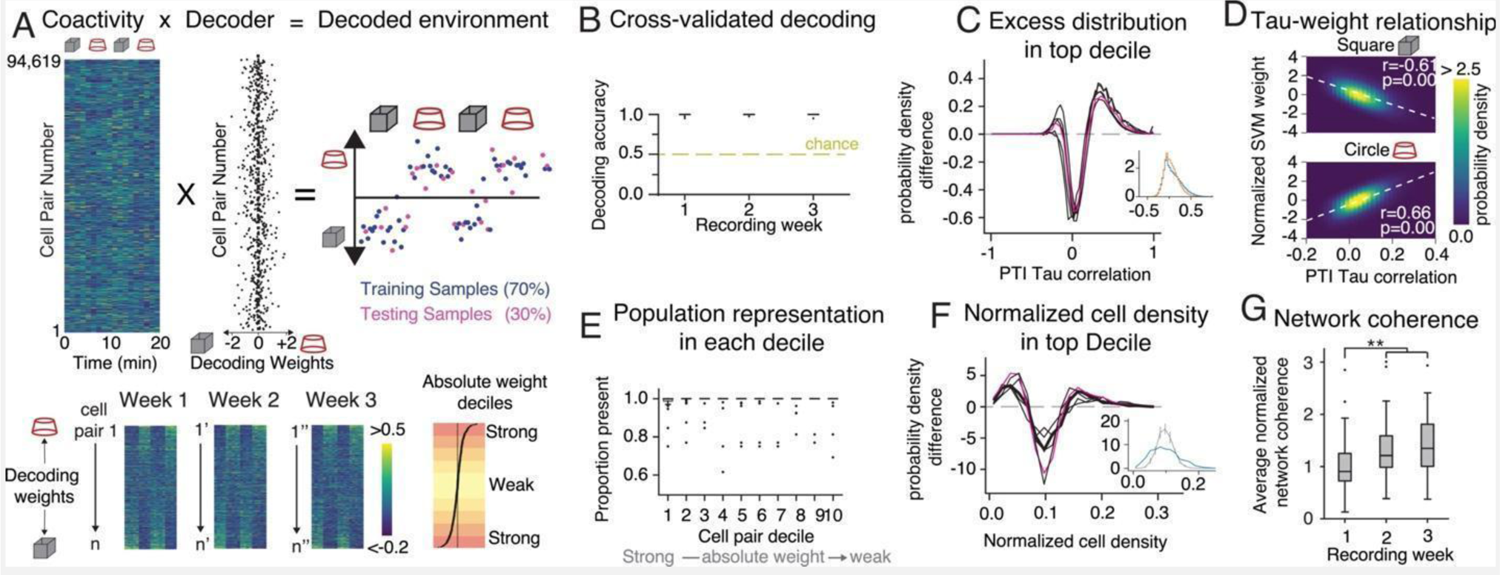
Decoding environments from PTI coactivity. (A) Illustration of SVM decoding from PTI coactivity; (top) PTI coactivity, computed every 60 s, is projected with SVM weights onto a single dimension that separates the two environments. (Bottom) Cell-pair coactivity from an example ensemble ordered by SVM weight each week; weights’ absolute value, indicative of separation strength, are classified by decile. (B) Cross-validated decoding accuracy is very high for all ensembles. (C) Difference between overall and first decile distribution of cell pair correlations (example distributions in inset and shown in purple in main figure) indicates importance of coactive and anti-coactive cell pairs, confirmed by (D) 2-D distribution of SVM weights against PTI coactivity. (E) Proportion of individual cells present in at least one cell pair in each SVM-ordered cell pair decile. (F) Difference between random and first decile, normalized distributions of individual cell participation (example distributions in inset and shown in purple in main figure) indicates overrepresentation of some cell in the first decile. (G) PTI network consistency normalized by the standard deviation of the cell-pair randomized distribution (Week: F_2,60_ = 15.76, p=10^-6^, η^2^ = 0.094; Dunnett’s tests: week 0 < weeks 2, 3).

### The environment-discriminating subset of CA1 cells

It is commonplace to describe hippocampal population activity patterns with activity vectors. These are distinct between the two environments, yet this difference is remarkably small (Fig. S7), consistent with the notion that the activity of only a minority of individual cells contribute to discriminating the environments. Because only a quarter of CA1 cells are place cells, and because only place cells change their position tuning and the firing of even fewer cells turns on or off across environments, this small change in population activity across environments might be expected if place cells drive the discrimination. Consequently, we ask what subset of the CA1 population contributes to discriminating the two environments. We trained an SVM decoder to project the data along a single composite dimension that separates the two environments. Using 1-s activity vectors recorded on the same day, the decoder correctly identifies the current environment well above chance; performance increases with the animal’s experience (Fig. 5A). After one week, decoding across days also performs above chance (Fig. 5B). Indeed, after one week, the SVM weights are stable across weeks; the weights obtained from the last day of the third week are strongly correlated to the weights obtained up to 9 days earlier, consistent with the expectation that SVM weight stability reflects learning, much like firing field stability has been traditionally interpreted (Fig. 5C). To evaluate which subset of cells contributes to discriminating the environments, we trained a decoder using select portions of the population to which the initial SVM decoder gave different weights. Decoders using just the 20% of cells with the largest weights perform indistinguishably from decoders that use the entire population. On the other hand, decoders using the 40% of cells with the smallest SVM weights perform close to chance (Fig. 5D). Decoders using cells with the 20% highest weights also display the largest increase in performance with experience, while for the bottom 60%, performance does not increase, consistent with the expectation that the largest SVM weights estimate the strength of spatial coding and learning, much like place field quality has been interpreted.

**Figure 5.**
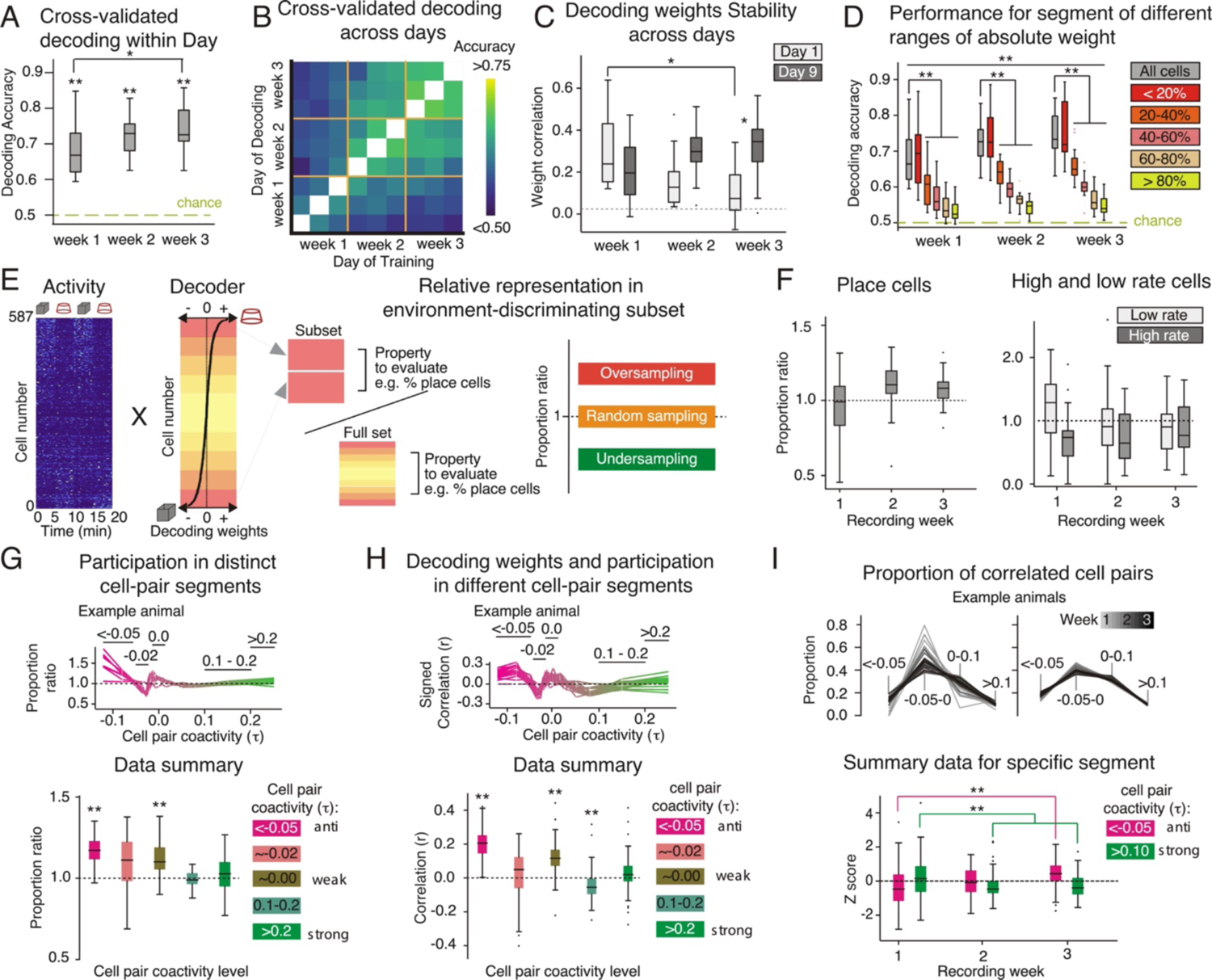
Decoding of 1-s activity vectors relies on a population subset and highlights the role of anti-coactive cell pairs. (A) Cross-validated decoding performance is above chance and increases each week (Week: F_2,12_ = 12.63, p=0.0011, η^2^ = 0.11; Dunnett’s tests: week 1 < week 3; post-hoc t tests vs. 0, **p<10^-7^) (B) Heatmap of cross-validated decoding performance across days. (C) Correlation of weights trained on day 1 and 9 with weights trained on other days, with weights from the same day excluded (No effect of Week: F_2,9_ = 2.11, p = 0.18, η^2^ = 0.011; no effect of Day: F_1,10_ = 1.71, p=0.22, η^2^ = 0.08; Interaction: F_2,9_ = 8.03, p = 0.01, η^2^ = 0.14; Dunnett’s tests: week 0 < week 2 on day 1; post-hoc t tests: day 1 vs day 9, *p < 0.017) (D) Cells are ordered by SVM absolute weights and separated into quintiles (20%) whose cross-validated performance is measured independently (Quintile: F_4,65_ = 38.58, p = 10^-16^, η^2^ = 0.65; Week: F_2,64_ = 29.86, p = 10^-10^, η^2^ = 0.022; Interaction: F_8,89.135_ = 3.69, p = 10^-4^, η^2^ = 0.026; post-hoc t tests: All Cells vs Quintiles, **p ≤ 0.001; Dunnett’s tests: week 0 < week 3 for quintile 1 and 2). (E left-right) Illustration of activity vectors with cells ordered by SVM weights. Weights’ absolute value, indicative of separation strength, are classified by quintile subsets. Тhe properties of the top quintile (subset), the environment-discriminating subset, are compared to the properties of the population (full set) to compute a proportion ratio that evaluates how cells from this subset differ from the population. (F, left) Ratio of proportion of cells classified as a place cell in either environment in the top quintile versus the overall population (No effect of Week: F_2,12_ = 3.58, p = 0.061, η^2^ = 0.13; t test vs. unity (1): t_48_ = 1.38, p = 0.18). (right) Ratio of proportion of cells with high or low rate in top quintile versus overall population (High Rate: No effect of Week: F_2,12_ = 0.65, p = 0.54, η^2^ = 0.015; t test vs. unity (1): t_48_ = 3.19, p = 0.0025; Low Rate: No effect of Week: F_2,12_ = 3.47, p = 0.065, η^2^ = 0.064; t test vs. 1: t_48_ = −0.07, p = 0.95). (G) Ratio of average participation in cell pairs of different coactivity levels for top quintile versus overall population (top) in an example animal and (bottom) for specific binned levels of coactivity (Coactivity Level: F_4,65_ = 5.71, p = 10^-4^, η^2^ = 0.094; No effect of Week: F_2,65_ = 0.19, p = 0.83, η^2^ < 10^-3^; No Interaction: F_8,128_ = 1.14, p = 0.34, η^2^ = 0.031; post-hoc t tests vs. 1, **p ≤ 10^-6^). (H) Average correlation between SVM weight and participation in cell pairs of different coactivity levels (top) in an example animal and (bottom) for specific coactivity levels (Coactivity Level: F_4,135_ = 55.53, p = 10^-27^, η^2^ = 0.43; No effect of Week: F_2,134_ = 1.57, p = 0.21, η^2^ = 0.0012; No Interaction: F_8,189.12_ = 1.82, p = 0.076, η^2^ = 0.018; post-hoc t tests vs. 0, **p ≤ 10^-7^). (I) Proportion of cell pairs with different coactivity levels across days (top) in two example animals and (bottom) for cofiring and anti-cofiring cell pairs (No effect of Coactivity Level: F_i,i22_ = 0.74, p = 0.39, η^2^ = 0.0043; Week: F_2,121_ = 3.91, p = 0.023, η^2^ = 0.0075; Interaction: F_2,121_ = 10.12, p = 10^-5^, η^2^ = 0.07; Dunnett’s tests vs week 1, **p ≤ 0.009).

We call this 20% of cells the “environment-discriminating subset” and evaluated their properties by computing whether the properties of cells in the subset are more or less prevalent than in the entire population (Fig. 5E). The place cells, strongly active cells, and weakly active cells are not more likely than chance to be in the environment-discriminating subset (Fig. 5F). We then evaluated whether the environment-discriminating subset is comprised of cells that tend to be coactive with other cells, or alternatively, tend to be anti-coactive with other cells. Only the anti-coactive cells are more likely than chance to be part of the environment-discriminating subset (Fig. 5G). Furthermore, the SVM decoding weights are predicted by the number of negatively correlated cell pairs to which a cell belongs. By contrast, participation in positively correlated pairs is not related to either SVM weights or being in the environment-discriminating subset (Fig. 5H). Consistent with anti-coactivity being an important feature for discriminating the two environments, we observe an increase in the number of negatively correlated cell pairs and a decrease in the number of positively correlated cell pairs, as mice learn to discriminate the two environments (Fig. 5I). This could be visualized by computing each cell’s anti-cofiring power as the proportion of the total significant negative correlations (τ ≤ −0.05) to which the cell contributes. Figure 6 shows ensembles recorded in weeks 1 and 3 as square matrices of the cell pair correlations. In rows 2 and 3 the matrices are organized according to increasing anti-cofiring power determined in either the cylinder or square environment. The anti-cofiring subset appears more stable after experience, consistent with learning (Fig. 5I). The most anti-cofiring cells can even be anti-correlated with each other, and the anti-cofiring subset is environment specific, with up to 40% overlap between the anti-cofiring subset of cells in the same environment on day 9 (Fig. 6C).

**Fig. 6.**
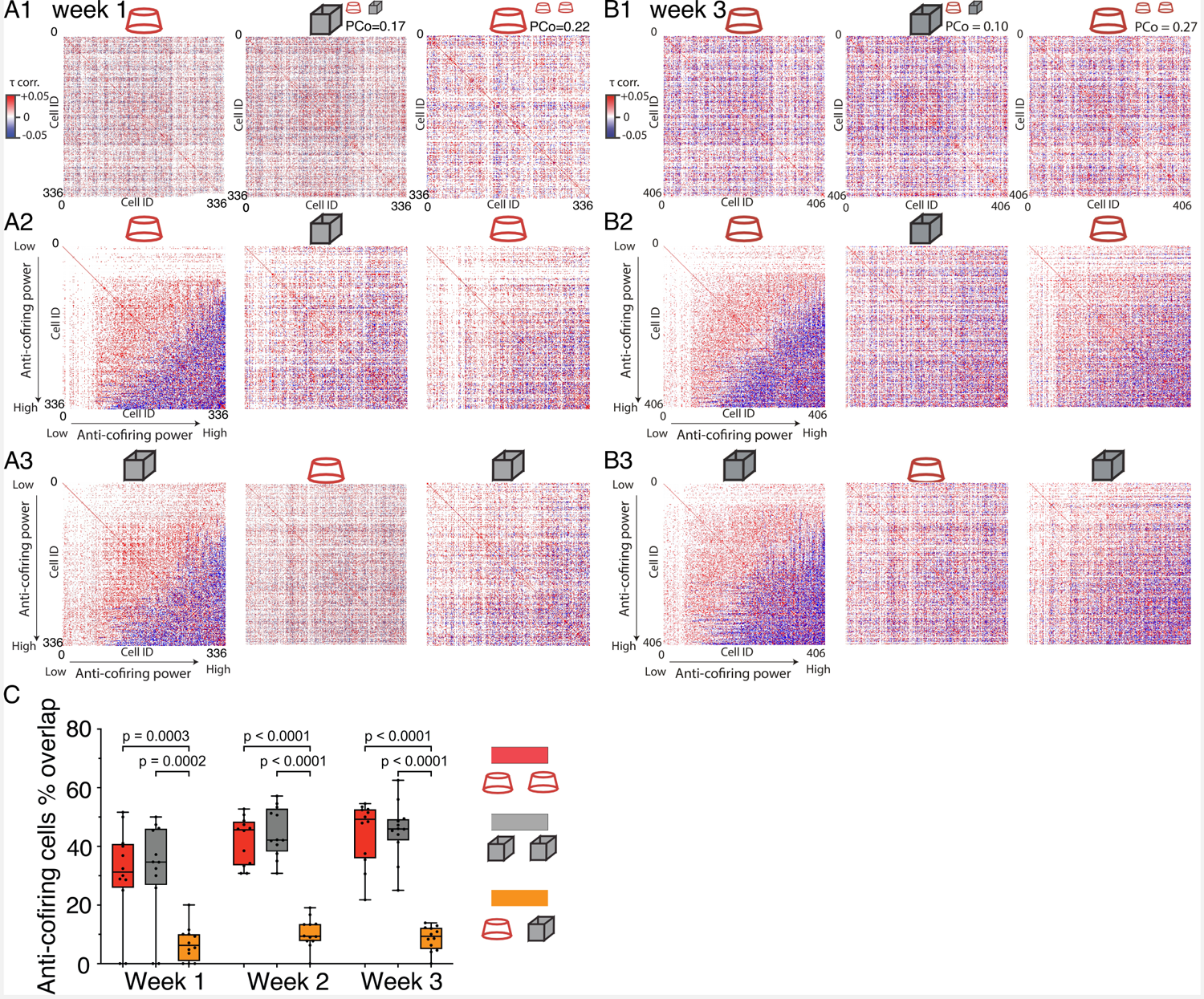
Anti-cofiring cells. Correlation matrices depicting all the pairwise correlations (τ) observed during three recordings in A) week 1 (day 1) and B) week 3 (day 9) from the same mouse. Statistically significant positive (red) and negative (blue) correlations are indicated. A1,B1) The cell ID order is arbitrary. A2,B2) The cell order is sorted according to anti-cofiring power in the first cylinder recording. A3,B3) The cell order is sorted according to anti-cofiring power in the first box recording. C) The percentage of cells that qualify as anti-cofiring in two recordings, as a function of week.

### Planar manifold topology of ensemble firing distinguishes environments

Place fields on a surface are planar, making it straightforward to imagine that place cell ensemble activity generates representations with 2-D structure that can differentially represent distinct 2-D environments. What is the dimensionality of the n-cell dimensions of ensemble activity in which coactivity relationships differentially represent environments? To evaluate the representational topology of the data we used persistent homology and computed Betti number barcodes (Ghrist, 2008) which indicate that the topology of 1-s CA1 ensemble activity patterns has one component and no holes. We assumed this might approximate a 2-D surface, and asked whether an unsupervised, non-linear dimensionality reduction algorithm can readily distinguish environments on a 2-D manifold (Fig. 7A; Video S1). The IsoMap algorithm uses the neighborliness amongst activity vectors to non-linearly project the data into an optimal low-dimensional subspace (Balasubramanian and Schwartz, 2002), so we compared it with PCA, a linear transformation of the data (see Video S2 to gain intuition). We also used the replica mean field theory of manifolds to characterize neural manifolds directly from high-dimensional ensemble activity vectors in terms of three geometric measures: capacity, extent, and dimensionality, without any assumptions on the low-dimensional geometry or topology of the data (Chung et al., 2018a). We first applied both the IsoMap and PCA algorithms to 1-s activity vectors recorded in the cylinder, box, and home cage environments. IsoMap distinguishes the ensemble activity vectors recorded in the three environments when the 1-s ensemble activity vectors are projected onto the two main IsoMap dimensions (Fig. 7B, right). The environments are harder to discriminate when the identical activity vectors are projected onto the two main PCA dimensions (Fig. 7B, left). These impressions are quantified using explained variance. The first 10 IsoMap dimensions explain ∼70% of the variance, compared to ∼20% with the first 10 PCA components (Fig. 7C). To quantify environment discrimination, we calculated for each recording the average projection onto the two main dimensions of IsoMap or PCA (Fig. 7D). The difference between projections from the <u>same</u> environments was conservatively normalized by the difference between projections from <u>different</u> environments (Fig. S8). This discrimination ratio will decrease from 1, when all environments are similarly represented, to zero, when the same environments are identically encoded, and different environments are well discriminated. Using IsoMap, this normalized vector difference decreases from 1 to about 0.3 with the animal’s experience (Fig. 7D). This contrasts with the modest decrease of the discrimination ratio when the activity vectors are projected onto the PCA dimensions, which stays above 0.8 on average. The discrimination ratio also only decreased modestly when it was computed on the raw activity vectors without any dimensionality reduction (Fig. 7D). Finally, to elucidate whether the contribution of coactivity is necessary and sufficient for the IsoMap discrimination of the environments, we recomputed the activity vectors after systematically removing or including only the top 5, 10, 25, and 50% of cells that participated most in the coactive and anti-coactive cell pairs. The surviving ensemble activity vectors were then projected on the two main IsoMap dimensions. Removal of the most coactive or anti-coactive cells selectively degraded the IsoMap discrimination (Fig. 7E), and inclusion of only the most coactive and anti-coactive cells was uniquely sufficient for the discrimination (Fig. 7F). These effects were stronger for the anti-coactive cells.

**Figure 7.**
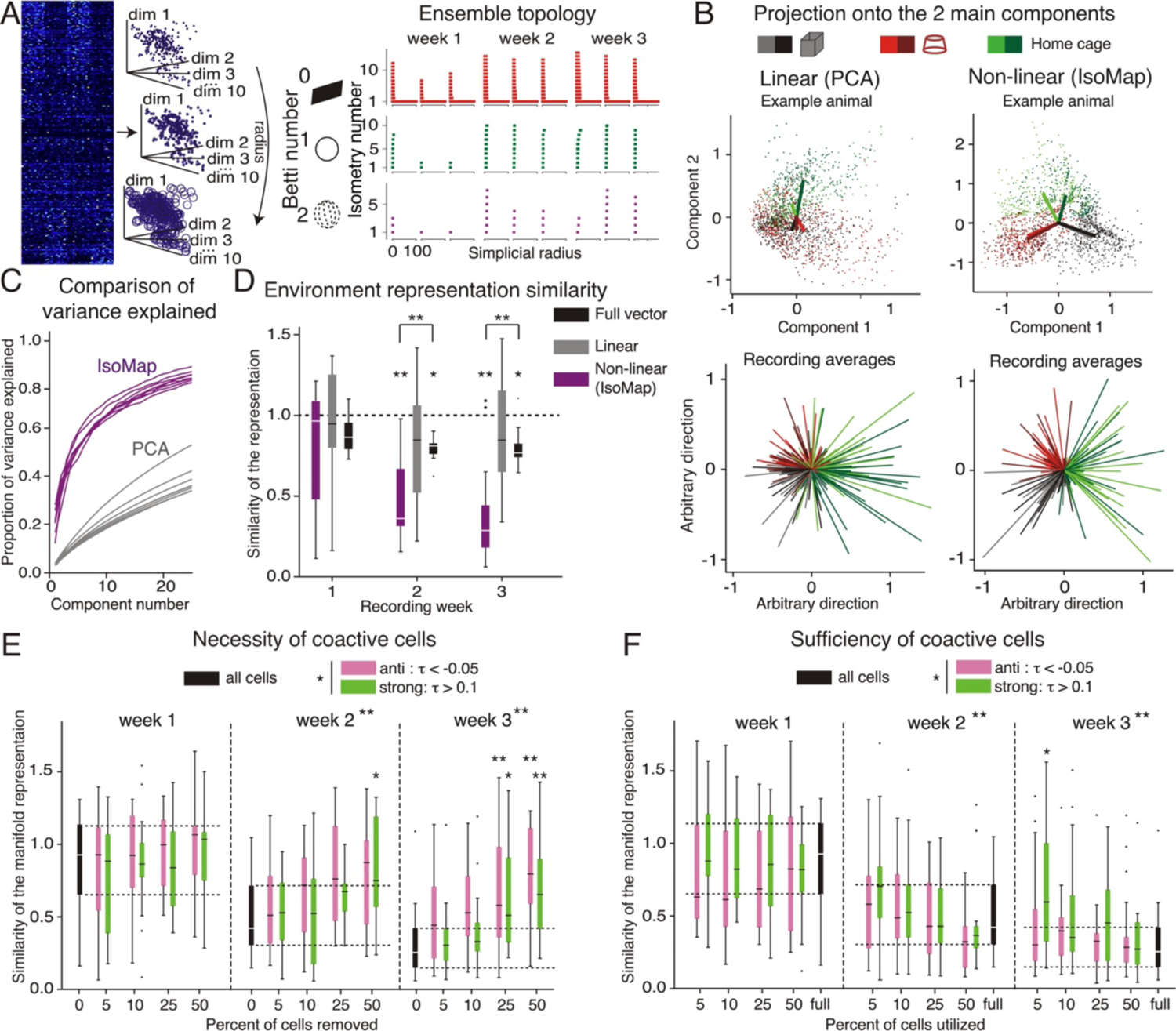
Non-Linear embedding distinguishes between environments reliant on acti-coactive cell pairs. (A, right) Illustration of the algorithm used to determine the Betti numbers that describe the topology of ensemble activity. (left) Activity vectors are projected onto a 10-dimensional space using IsoMap and points are connected to neighbors with increasing radius of each point. Isometries are determined at each radius. (right) Example of Betti barcode showing that only one isometry of Betti number 0 persists, indicating that the population activity appears constrained to a single connected subspace. (B) Activity vectors projected onto a 2-D subspace using a linear (PCA) or nonlinear (IsoMap) dimensionality reduction algorithm. In each recording the average 2-D vector is computed as shown (top) in an example animal with individual points, projections of individual 1-s activity vectors, and (bottom) for all recordings and animals. Repetitions of the same environments on the same day are given distinct color shades. (C) Projections onto 3-D and higher dimension subspaces were also performed. The proportion of variance explained as a factor of the number of dimensions for PCA and IsoMap is given, indicating that the non-linear 2-D IsoMap subspace explains substantially more variance than the linear PCA projections. (D) Ratio of the difference between average vector from similar environment and different environments using IsoMap, PCA, or the full, non-dimensionally reduced, activity vectors (Method: F_2,39_ = 10.38, p = 10^-4^, η^2^ = 0.19; Week: F_2,38_ = 6.67, p = 0.003, η^2^ = 0.076; Interaction: F_4,44.58_ = 2.96, p = 0.03; post-hoc Dunnett’s tests: IsoMap < Full Vector, **p ≤ 0.046; post-hoc Dunnett’s tests: week 1 > weeks 2, 3, *p < 0.03, **p < 0.01). (E) Similar ratio after removing different percentages of the population as a function of their contribution to cell pairs of different coactivity levels (Coactivity Level: F_1,99_ = 6.06, p = 0.016, η^2^ = 0.012; Percentage: F_1,99_ = 17.86, p = 10^-5^, η^2^ = 0.078; Week: F_2,98_ = 22.08, p = 10^-8^, η^2^ = 0.13; no Interaction, all p’s > 0.18; post-hoc: Anti > Strong t_380.93_ = 2.11, p = 0.036; post-hoc Dunnett’s tests: week 1 > weeks 2, 3, **p ≤ 10^-7^; post-hoc t tests vs. All Cells, *p’s < 0.05, **p’s < 0.01). (F) Similar ratio using only percentages of the population as a function of their contribution to cell pairs of different coactivity levels (Coactivity Level: F_1,99_ = 5.27, p = 0.24, η^2^ = 0.014; Percentage: F_1_,_99_ = 4.71, p = 0.032, η^2^ = 0.025; Week: F_2,98_ = 13.00, p = 10^-6^, η^2^ = 0.17; Week x Percentage x Coactivity Levels: F_2,98_ = 3.10, p = 0.049, η^2^ = 0.0055; no other significant interaction, all p’s > 0.12; post-hoc: Anti < Strong t_380.40_ = 2.33, p = 0.020; post-hoc Dunnett’s test: week 1 > weeks 2, 3, **p < 0.01; post-hoc t tests vs. All Cells, *p’s < 0.05).

These findings motivate a novel “reregistration” hypothesis that CA1 ensemble coactivity can discriminatively represent distinct environments through a relatively conserved neuronal ensemble activity pattern that is constrained on a non-linear surface that is at once conserved across environments, and distinctively registered to each environment as a result of the experience-dependent development of anti-coactive neural discharge (Fig. 1B; 7A).

Importantly, the hypothesis asserts that environment-specific differences in cofiring relationships amongst the cells as shown in Figs. 3,6, are largely due to the anti-cofiring cells, that this subset of cells is responsible for the differences between environment-specific manifolds of neuronal activity, and that the topology of the environment-specific manifolds is invariant (Fig. 8B). Anti-cofiring power was computed for each cell in an ensemble as the proportion of its ensemble cell pairs for which τ < −0.05 (Fig. 8C). Both cells of an anti-cofiring pair tended to be active rather than one cell active and the other silent in a particular environment (Fig. S8).

**Fig. 8.**
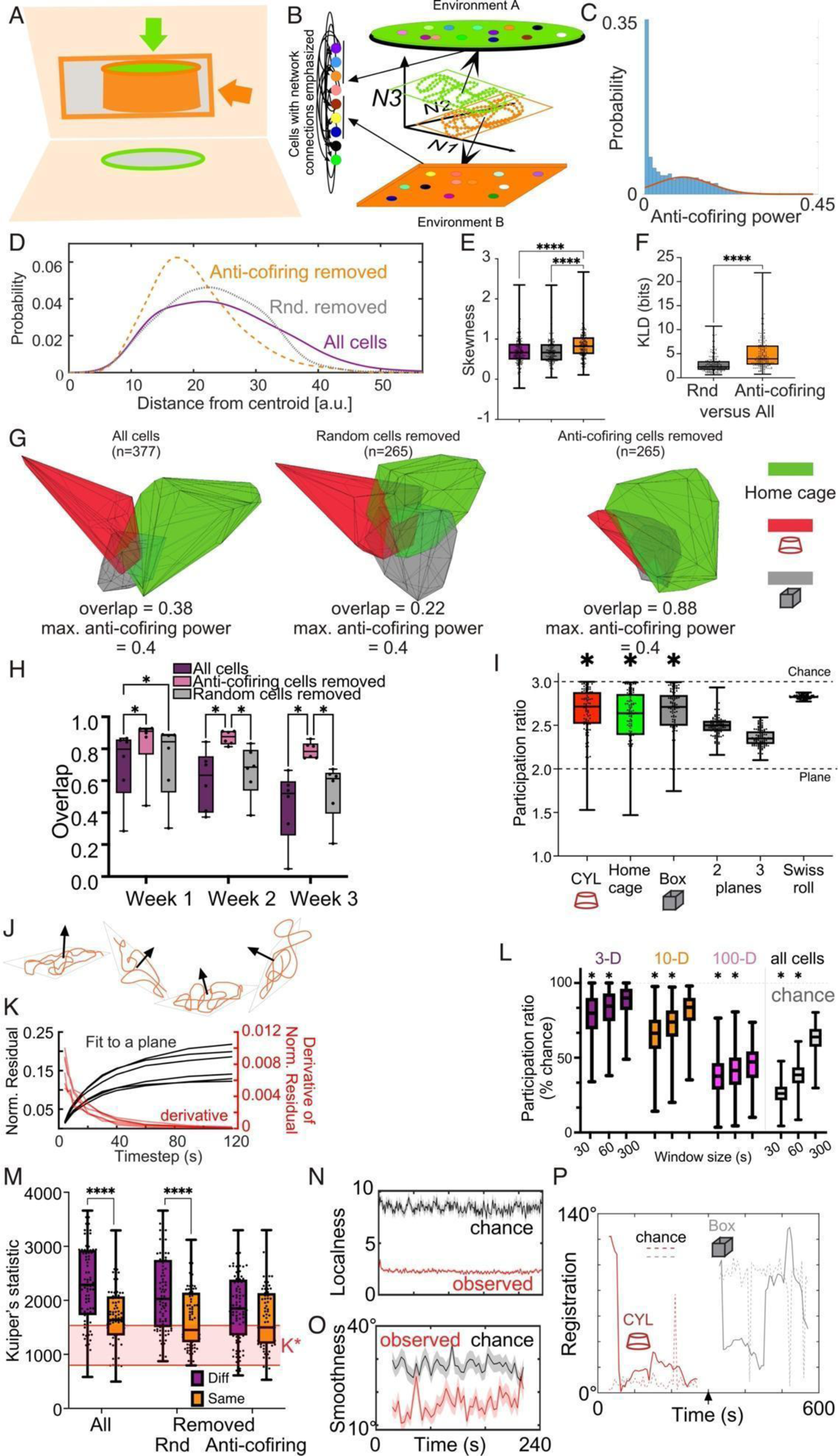
Reregistration, reconceptualizing how hippocampus represents multiple environments. (A) Schematic illustrating that projections of an invariant object (a solid cylinder) at distinct angles can generate distinct representations (circle or rectangle). (B) The reregistration hypothesis illustrated. Intrinsically-organized neuronal network activity organizes as smooth trajectories on a low-dimensional surface. A low-dimensional (2-D) manifold (green and orange rectangles) is depicted in the 3-D subspace. Environment-specific representations result from projecting the environment onto the manifold, which does not reorganize cofiring relationships, but reorganizes place cell firing fields (color ellipses). Environment-specific registrations of the manifold with external cues (different angles in the 3-D subspace) result from taking perceptual fixes that orient manifold activity with respect to external cues. (C) Distribution (blue) with Gaussian fit of the anti-cofiring tail (orange) of coactivity power computed for cells from each recording in the dataset. (D) A day-7 recording example histogram of 300 1-s activity vector distances from the centroid after projecting a 377-cell (All) ensemble into a 60-D IsoMap subspace. The 112 (30%) most anti-cofiring and an equivalent random set of cells were removed, and the remaining 265-cell ensembles were projected into a 60-D IsoMap subspace. The histogram comparisons suggest that the anti-cofiring cells are disproportionally far from the centroid. The example is from the second recording in the rectangular environment during the third week, for mouse M39. Summary quantification of (E) Skewness and (F) Kullback-Liebler divergence of the 3 histograms from each of all recordings (n = 200) confirm anti-cofiring cells are distinctive and significantly distant from the centroid. (G) Visualization as a volume, after projecting the 1-s ensemble activity vectors from panel D into a 3-D subspace defined by the first 3 IsoMap components. The component coordinate system was determined from two 5-min recordings of the ensemble population in the home cage, box, and cylinder. The volumes occupied by activity vectors were compared for all cells (left), a subset of randomly selected cells (middle) or after removing the most anti-cofiring cells (right). Removing the 30% most anti-cofiring cells collapsed the distinction between the cylinder and box volumes. (H) Summary data from all ensembles recorded across the 3 weeks illustrates that the distinction between the environment-specific manifold volumes increases with experience, and that the anti-cofiring subset of cells are crucial for the distinction (two-way Ensemble x Week ANOVA, Ensemble: F_2,45_= 14.04, p = 10^-5^; Week: F_2,45_= 7.33, p = 0.002; Interaction F_4,45_= 0.32, p = 0.9; Friedman statistic = 30.79, Anti-cofiring removed > All cells = Randomly removed). (I) Participation ratios (PR) estimate the dimensionality of the ensemble activity projected into the 3-D IsoMap subspace for all the recordings. The dimensionality is significantly different than chance (PR=3) and indistinguishable from planes with multiple shifts as well as a non-linear Swiss Roll (Kruskal-Wallis statistic = 138.2, p = 0). (J) Schematic illustrating the hypothesis that ensemble activity in the 3-D IsoMap subspace is organized along planar trajectories that episodically shift orientation in the subspace. (K) Residuals and derivative from best fit planes for each mouse as a function of the data timestep used to estimate the duration ensemble activity is constrained to a plane in the 3-D IsoMap subspace. (L) Participation ratio data dimensionality estimates by projecting different windows of data into IsoMap subspaces of different dimensionality. (M) Kuiper’s statistics compared the set of average angles of the best fit planes in the 3-D IsoMap subspace for each of the daily recordings in the same or different box and cylinder environments. (N) Summary measures of the localness and (O) smoothness timeseries of the 1-s ensemble activity trajectories in the 3-D subspace compared to chance and (P) an example timeseries (4-s timestep) of the angular registrations of the best fit plane to 60-s of data along the ensemble trajectory through the 3-D IsoMap subspace.

We then investigated whether the anti-coactive cells cause the ensemble activity to be distinctive by projecting the data into a 60-D IsoMap subspace. The 60-D choice facilitated statistical comparisons by allowing all ensembles to be assessed in a subspace of the same dimension that is sufficient to account for most of the variance in activity vectors (Fig. 7C). The 60-D centroid of the 1-s activity vectors from each recording was computed and the distribution of the Euclidean distance from the centroid for each activity vector was examined. Fig. 8D, the example of a 377-cell ensemble recorded on day 7, illustrates the distribution has a long tail of ‘outlier’ vectors that are far from the centroid. We tested if the most anti-coactive cells disproportionately contribute to the outliers by removing the 30% (112) most anti-coactive cells from the ensemble and as a control, removed an equivalent random sample of cells then recomputed the histogram. Removing the anti-coactive cells reduced the number of ‘outliers’ and strongly shifted the distribution leftward more than the random removal. This impression was quantified by computing the skewness of the three distributions for each recording across the mice (Fig. 8E; Friedman statistic = 69.12, p < 0.0001; 3 distributions, each with 200 samples; anti-coactive cells removed > all cells = random cells removed, p’s < 0.0001). Using the Kullback-Liebler divergence (KLD) we then compared how different the histograms of the distances from the centroid became after removing the anti-coactive and random cells.

Consistent with the example in panel D, removing anti-coactive cells increased the difference from the histogram of the full ensemble far more than removing random cells (Fig. 8F; Wilcoxon W=18260, p < 0.0001). These findings from the 60-D IsoMap subspace suggest that anti-coactive cells cause distinctive activity vectors. This effect is confirmed using the replica mean field theory of manifolds that we used to evaluate the relative change of dimensionality and extent measures between two scenarios (see Methods): removing anti-cofiring cells versus removing the same number of random cells. The relative change in the extent of the manifold (5.83± 0.76) is far larger than the increase in dimensionality (3.27 ± 0.56; paired t test t_49_ = 6.14, p = 10^-7^), hinting that the contributions of the anti-coactive cells are closely linked to the variability of the anchor points on the margin of the manifolds, thus changing the extents of these manifolds (Fig. S10).

Next, to visualize and test the possibility that the anti-cofiring cells are responsible for the distinction between the environmental representations, we projected the activity vectors from the pair of cylinder and box recordings on a particular day into a 3-D IsoMap subspace. The three principal IsoMap components were determined by concatenation of the home cage, box, and cylinder recordings of 1-s activity vectors, and projected into the same 3-D sub-space. Figure 8G shows the volumes occupied by the 600 (2 x 5 min) 1-s ensemble activity vectors from the 377-cell ensemble recorded on day 5 in the home cage, cylinder, and box for mouse M39. The overlap of the cylinder and box activity is modest (0.38), consistent with ensemble activity being sufficiently distinctive to discriminate the two environments (Fig. 7). Removing a random 30% of cells did not increase the overlap (0.22) whereas removing 30% of the most anti-coactive cells reduced the outliers (by inspection), did not remove enough cells to change the maximum anti-cofiring power, and importantly, collapsed the distinction between the cylinder and box activity patterns (0.88). We repeated this analysis for the second or third recording during each week to determine whether the ability of the anti-cofiring cells to distinguish the cylinder and box ensembles developed with experience. Figure 7H illustrates that during the first week, the cylinder and box activity vectors occupied a largely overlapping subspace but with experience, from week 2, they became more distinct. Importantly, removing the anti-cofiring cells, which reduced the maximum anti-cofiring power from 0.20±0.013 to 0.15±0.013, reduced the distinction between the two environments, but not when an equivalent subset of cells was randomly removed (maximum anti-cofiring power 0.19±0.013). The reduced distinctiveness is also observed in the capacity metric of the replica mean field theory of manifolds, the high-dimensional measure of overlap that we used IsoMap dimensionality reduction to estimate (see Methods). Relative to removing the same number of random cells, removing anti-coactive cells decreased capacity (−5.13 ± 0.74; Fig. S10). This pattern is predicted by the reregistration hypothesis.

### Multistable manifold representations of environments

Projecting the activity vectors into the volume of the first three IsoMap components, hinted they may organize on 2-D surfaces (see Videos S1,S3), which is corroborated by computing participation ratios (PR) to estimate the dimensionality (PR = 2.65 ± 0.30, n = 258) (Gao et al., 2017; Recanatesi et al., 2019). These ratios were significantly lower than chance (PR = 3), but indistinguishable from a non-linear Swiss roll surface, or 2-D planes that change orientation within the 3-D space (Fig. 8I).

We’ve previously observed that internally-organized entorhinal and hippocampal spatial representations are multistable, they spontaneously change registration to the environment on the time scale of ∼10 seconds during free navigation (Chung et al., 2021; Dvorak et al., 2021; Dvorak et al., 2018; Fenton et al., 2010; Kelemen and Fenton, 2016; Park et al., 2019; Talbot et al., 2018). Accordingly, we considered the possibility that CA1 activity is multistable, in that it intermittently registers to distinct environmental features, which would change the orientation of a fundamentally lower-dimensional (2-D) surface within the higher-dimensional (3-D) IsoMap space, as cartooned in Fig. 8J.

Consistent with the multistable conjecture, 30-60 s of activity vectors are well fit to a plane in the 3-D IsoMap subspace (Fig. 8K). We animated the activity vector timeseries projected into the IsoMap subspace (Video S3), which upon inspection shows the high-dimensional ensemble activity takes non-random, smooth trajectories through a 2-D subspace that intermittently changes orientation. Analysis of participation ratios computed on 30, 60, and 300-s time windows (Fig. S9) indicate that the dimensionality is significantly lower for the shorter time windows (Fig. 8L) suggesting that the hypothesized multistability occurs on a time scale of ∼60 seconds.

We proceeded by computing Kuiper’s statistic to measure whether the set of 60-s planar angles differed for daily recordings in the same and different box and cylinder environments. Fig. 8M shows that the angles were more similar in two recordings in the same environment than they were in different environments, consistent with Figure 7, and indicating that manifold orientation in the subspace can distinguish the two environments. Importantly, this distinction between recordings in the two environments is lost when anti-coactive, but not random cells are removed (Two-way Environment x Ensemble ANOVA, Environment: F_1,534_ = 56.87, p = 10^-13^, Ensemble: F_2,534_ = 6.77, p = 0.001; Interaction F_2,534_ = 5.06, p = 0.007; post-hoc Tukey: Different environments: Anti-coactive removed < All cells = Randomly removed p’s < 0.02; Same environments: Anti-coactive removed = All cells = Randomly removed, p’s > 0.98). We quantified the localness (Fig. 8N; 2.26± 0.01 vs. chance: 8.44± 0.04, Wilcoxon W = 8.77×10^8^, p < 10^-4^), and smoothness (Fig. O; 16.5°± 0.4° vs. chance: 28.0°± 0.39°, Wilcoxon W = 1.68×10^7^, p < 10^-4^) of the ensemble trajectory in the IsoMap subspace, which are distinct from chance and together indicate that ensemble activity projected into the three best components of the IsoMap subspace is largely constrained to a 2-D plane for several 10’s of seconds, and then within 1-2 seconds, the plane reorients in the IsoMap subspace (Fig. 8P).

Analogous to firing rate maps, the spatial distribution of each of the three principal IsoMap components of the ensemble activity vector projections were computed to investigate the possibility of time-averaged spatial tuning (Fig. S11A). By inspection, no obvious patterns were recognized, but they were also not random because within week 2 recordings (days 4,5, or 6), the patterns across the same environments were significantly correlated for 4/6 mice (10/36 correlations were significant), and also across the different environments for 5/6 mice (7/72 correlations were significant). Finally, we examined the activity vectors around the times of the reorientations but did not recognize any relationships (r’s < 0.14, p’s > 0.2; Fig. S11B).

## Discussion

### Place field and coactivity-based neural codes for space

We find that the coactivity relationships amongst CA1 principal cells discriminate environments and locations, and the cofiring stabilizes with experience (Fig. 3, S5), identifying a learned neural code that can be independent of the place field-based code conventionally used to understand the hippocampal role in spatial cognition. Instead of single cell properties, our analyses focused on the set of pairwise activity correlations that approximate the higher-order activity correlations that may define informative neural population activity patterns (Fig. S3; Ebitz and Hayden, 2021; Jazayeri and Afraz, 2017; Schneidman et al., 2006). We took advantage of the miniature microscope technology to monitor the activity and positions of large numbers of the same cells over weeks (Fig. 2), an ability which was not available with the electrophysiological techniques that established the importance of place fields and remapping for the representation of space. There are of course limitations to this technology that impact the present study. The timescale of analysis was restricted to 1-s as processes faster than ∼300 ms are precluded by the slower time scale of GCaMP6f transients. Nonetheless, 1-s is the behavioral timescale of CA1 synaptic plasticity mechanisms (Bittner et al., 2017; Milstein et al., 2021), and sufficient time resolution for computing the session-long cofiring relationships because the statistics of cofiring relationships measured at 25 and 40 ms are equally well captured by the statistics of 1-5 s cofiring relationships (Olypher et al., 2006; Park et al., 2019).

We restricted analyses to anatomically separated cell pairs to avoid spuriously high activity correlations, but this prevents analysis of the fine-scale topographical organization of the neural circuit (Hampson et al., 1999; Pavlides et al., 2019). Place cell stability was variable with some cells unstable (similarity ∼ 0) and others highly stable (similarity > 0.5). Stability increased after the first 3 days (30-min total) of exposure (Fig. 2F,G) reaching a magnitude that is typical of mouse place cells recorded with tetrodes during purposeful behavior (Kentros et al., 1998; Muzzio et al., 2009). Network consistency, which estimates the alignment of higher-order activity correlations, also increases with experience and with coactivity discrimination of environments and locations, further highlighting the importance of coactivity (Fig. 4, Fig. S5), consistent with the modeling results (Figs. 1, S2), and reports that CA1 place field plasticity relies more on population cofiring statistics than on the activity of individual cell pairs (Milstein et al., 2021). Coactivity relationships were scale-free (Fig. S3) and dominated by a minority of anti-coactive cells for discriminating environments (Figs. 4,5,7,8), features that describe the behavior of flocks and their interactions with external features (Hemelrijk and Hildenbrandt, 2011, 2012; Reynolds, 1987). Support vector machine decoding objectively measured the globally optimal contribution of each cell or cell pair to distinguishing the two environments (Figs. 4,5) and locations (Fig. S5). While this was effective on a 1-s timescale, it only demonstrates the availability of discriminative information, rather than how information is encoded and extracted by neural circuits, not unlike how the finding of place fields has not determined how the hippocampus uses them. The findings identify the importance of coactivity for representing environments, however these data in no way preclude place field-based codes for representing locations; decoding location with place cells was certainly better than without place cells (Fig. S5). Rather, both coding schemes may operate in parallel, and more likely together. We note that the possibility of having multiple firing fields in larger environments (Fenton et al., 2008; Harland et al., 2021; Muller and Kubie, 1987; Park et al., 2011) can degrade decoding of environments (Fig. S1) and the literature reports diverse additional phenomena that may also complicate such dedicated place codes. These phenomena include fundamental features of these cells such as firing-rate overdispersion (Fenton et al., 2010; Fenton and Muller, 1998; Jackson and Redish, 2007; Nagele et al., 2020; Poucet et al., 2012), mixed tuning to extra-positional variables (Fusi et al., 2016; Hardcastle et al., 2017; Stefanini et al., 2020), theta-phase temporal coding (Buzsaki and Chrobak, 1995; Harris et al., 2003; Hirase et al., 1999; Huxter et al., 2003), and firing-field instability (Fig. 2; Muzzio, 2018; Ziv et al., 2013), which should be properly evaluated by simulation studies (Kang et al., 2021). We also relied on the IsoMap algorithm to discover that the somewhat variable ensemble activity was constrained to low-dimensional subspaces where distinct environment-specific patterns relied on coactivity (Fig. 7, 8). We analyzed ensemble activity within abstract 60-D IsoMap subspaces that account for almost all the variance (Fig. 7C), as well as assuming a temporally-local 2-D organization within a 3-D IsoMap subspace to evaluate the multistability conjecture without assuming knowledge of the true geometry, or that the true dimensionality is 2-D; others have argued the neural topology reflects the spatial geometry of the environments from which place cells are recorded (Chen et al., 2014). Despite making such simplifying assumptions, our findings are robust; they hold when we applied the replica theory of mean field manifolds that measures rather than assumes the network geometry, without dimensionality reduction (Chung et al., 2018a). Taken together, these findings constrain the possible neural mechanisms for reading out the information represented in coactivity and point to synapsemble properties that have been proposed (Buzsaki, 2010; Buzsaki and Tingley, 2018) and are demonstrated, in artificial neural networks, to be effective mechanisms for transforming variable input activities to a reliable readout (Heeger and Mackey, 2019).

### The roles of feature tuning, coactivity, recurrence, and stability in the neural space code

Our experimental design (Fig. 2A) is the classic remapping experiment repeated across weeks, with recordings during all exposures to the environments. We replicated the fundamental findings that single cells change their place fields between environments and reinstate the place fields upon imminent return to an environment (Fig. 2F,H). While recurrence manifests as this place field stability and is standardly interpreted as memory persistence (Leutgeb et al., 2005c; Wills et al., 2005), it is hard to reconcile with the replicated finding that place fields are also unstable across weeks even in familiar environments (Fig. 2F,J; Lever et al., 2002; Ziv et al., 2013). This would require numerous (∼n) cell-specific rules to map the place fields from n cells in order to maintain, from one experience to the next, the coherent representation of the environment that a memory system requires (Lever et al., 2002; O’Keefe and Burgess, 1996). Furthermore, for such a scheme to be useful for discriminating environments, place cells should discharge predictably in their firing fields, but they do not (Fenton et al., 2010; Fenton and Muller, 1998; Jackson and Redish, 2007; Leutgeb et al., 2005b). In contrast, we identified recurrence of moment-to-moment coactivity relationships amongst CA1 cells, that the recurrence is modest (Fig. 3C), but sufficient for differentially representing environments as well as locations (Fig. 4B, S5).

Importantly, recent findings demonstrate that variable, distributed activity can organize within the stability of a manifold representation that explicitly does not rely on the reliability of either single cell or population activity, the way that conventional place field-based and activity vector-based spatial codes have been conceived (Figs. 6,7; Chaudhuri et al., 2019; Nieh et al., 2021; Rubin et al., 2019). Our cofiring analyses focus on the internal organization of neuronal activity, based on the firing of other neurons, rather than external variables like places, as demonstrated to be effective in hippocampus (Curto and Itskov, 2008; Keinath et al., 2022; Pettit et al., 2022) as well as MEC, a major input to hippocampus (Gardner et al., 2022; Low et al., 2022; Park et al., 2019; Yoon et al., 2013). The present findings offer an alternative concept for how hippocampal activity can be persistently informative without firing field stability (Fig. 2), or even population activity stability (Fig. 3; Video S2). A connectivity weight matrix, in other words the hypothesized synapsemble (Buzsaki, 2010; Buzsaki and Tingley, 2018), can transform the non-linear low-dimensional projections of neuronal activity (Fig. 7,8) to a steady readout (e.g., environment identity and location), so long as the connectivity matrix is invariant to shifts in the manifold axes, in which case the manifold would subserve memory persistence despite instability of neural activity, as demonstrated using artificial neural networks (Heeger and Mackey, 2019).

### Reregistration instead of remapping to represent spaces and memory

The remapping concept is founded on the notion that spatial and thus temporal firing relationships amongst hippocampal neurons change across environments and memories. This interpretation is based on the observation that firing fields change across, but not within environments, and the inference that ensemble temporal firing relationships change in correspondence (Colgin et al., 2008; Kubie et al., 2020; Muller and Kubie, 1987). As reviewed above, these assumptions and related inferences are violated (Fig. 2). Indeed, CA1 ensemble activity is only modestly different across environments (Fig. 3, Fig. 8) and within environments, can exhibit multistable dynamics that resemble remapping expectations (Dvorak et al., 2018; Fenton, 2015a; Jackson and Redish, 2007; Kao et al., 2017). Taken together, the data are consistent with the alternative concept of “reregistration.” Reregistration assumes CA1 activity is internally-organized in a relatively invariant manner such as on a low-dimensional manifold (Fig. 8A,B). Instead of external stimuli causing the rearrangement of neuronal activity, the internally-organized activity maintains but variably registers to external stimuli (Rule et al., 2019). This process may resemble fitting a model to observations (Fig. 8B). Cells that are most strongly influenced by currently salient stimuli are most likely to change their firing and because of the scale-free network correlations (Fig. S4) they influence and are influenced by the activity of all other cells. In flocks this allows the activity of most individuals to maintain locally while the entire flock changes globally (Hemelrijk and Hildenbrandt, 2011, 2012; Reynolds, 1987). The result of reregistration is that only a minority of cells change their activity distinctly from the activity of the other cells in the network, and according to present findings (Figs. 4-8), these we speculate are the anti-cofiring cells. The global network activity retains its low-dimensional organization but is registered (i.e. attached) uniquely to the environment in a manner that persists, especially we speculate, if synaptic weights are preferentially adjusted between the anti-cofiring cells and their environment-associated inputs. Because place cell firing is fundamentally multimodal, the activity only appears unimodal in small environments (Fenton et al., 2008; Harland et al., 2021), the ∼25% of CA1 cells that express firing fields change firing field locations with reregistration, although the internal organization of activity changes much less, as confirmed by simulations (Fig. 1, S2). Reconceptualizing remapping as reregistering identifies the importance of the anti-cofiring cell subset, not only as crucial for discriminating environments (Fig. 4-8), but also for orienting the relatively invariant manifold activity patterns within the neuronal activity subspace (Fig. 8).

CA1 activity has been recorded during explicit memory tasks to observe how place fields change with memory (Frank et al., 2004; Jeffery et al., 2003; Lenck-Santini et al., 2001). Although the field has struggled to establish a firm relationship to firing fields, memory correlates have been identified in cofiring relationships between excitatory as well as inhibitory cells in sleep and active behavior (Dvorak et al., 2021; O’Neill et al., 2008; Skaggs and McNaughton, 1996; van Dijk and Fenton, 2018). This is in line with the original conceptualization of remapping as a reorganization of the temporal discharge properties within an ensemble (Kubie et al., 2020; Kubie and Muller, 1991). Such changes in coactivity are consistent with synaptic plasticity studies that demonstrate balance (Okun and Lampl, 2008), bidirectionality (Milstein et al., 2021), and involvement of both excitatory and inhibitory cells (Basu et al., 2016; Caroni, 2015; Chung et al., 2021; Mongillo et al., 2018; Ruediger et al., 2011). We find that the strongly coactive and anti-coactive cell pairs are particularly discriminative at the 1-s timescale and, remarkably, anti-coactive cells are both more discriminative and more important for organizing the activity on a manifold (Fig. 6-8; Okun et al., 2015). This is consistent with the increase of interneuron-principal cell cofiring that has been observed at moments of memory discrimination and recollection (Dvorak et al., 2021; van Dijk and Fenton, 2018). Using PTI, we analyzed activity fluctuations around place tuning (Agarwal et al., 2014), in effect removing the place signal from the time series. We did this to compute cell-pair coactivity beyond cofiring due to place tuning and found that it improved the ability to discriminate environments (Fig. 4 and Suppl. Fig. 1), hinting at the possibility of relative independence of the place and environment codes. To be clear, place cells, their place tuning and their coactivity contribute to location-specific place coding more than non-place cells (Fig. S5) and no claim is made that place cells are irrelevant. We stress that our decoding of location and environment information from neural activity is simply to demonstrate the information is present in the data being decoded, and in no way do we suggest the decoding methods we have used are implemented by neural circuits. The focus of this work has been to examine the role of cofiring, which has been far less studied than the role of place fields, and both features of neural activity likely contribute to neural mechanisms of spatial cognition. There are already precedents of parallel codes for distinct spatial variables (Harris et al., 2003; Huxter et al., 2003; Meshulam et al., 2017; Poucet et al., 2012; Sarel et al., 2017; Stefanini et al., 2020; Tanaka et al., 2018). The coexistence of place field and coactivity codes for different features of space in hippocampal neural activity predicts that the explicit memory-related information in CA1 activity that has evaded the field’s efforts, would manifest in short-time scale coactivity, although in our view the manifold organization of observed ensemble coactivity relationships, replete with spatial modulation, is more likely the proper description of the hippocampal spatial code. We recognize that while we have identified a correspondence between patterns of neural coactivity and the ability to discriminate environments, we, like the rest of the field, have not determined what is actually represented in neural activity nor whether this correspondence causes the subject to understand its environments as distinct, or anything else for that matter. Nonetheless, our findings demonstrate the value of a conceptual shift towards a serious consideration of such vexing problems from the vantage of the collective and inherently temporally-structured behavior of neural activity (Brette, 2019; Buzsaki, 2019).

## Acknowledgements

Supported by NIH grant R01MH115304. We are grateful to Dr. John Kubie for valuable comments on an earlier draft of the manuscript, and Kathryn McClain for the initial network simulations that validated the network consistency measure.

## Author Contributions

A.A.F. and E.R.J.L. designed research; E.R.J.L., E.P. performed physiological research; J.H. and W.R., performed modeling research; S.C. provided analytical tools and knowhow; E.R.J.L., S. C.-S., and A.A.F. analyzed data; A.A.F. wrote the paper with contributions from all coauthors; A.A.F. supervised research.

## Methods

### Ethics Approval

All work with mice was approved by the New York University Animal Welfare Committee (UAWC) under Protocol ID: 17-1486.

### Virus injection and lens implant

To virally infect principal cells to express the fluorescent calcium indicator GCaMP6f, adult C57BL/6J mice (n=41) were anesthetized with Nembutal (i.p. 50mg/kg), one hour after receiving dexamethasone (s.c. 0.1mg/kg). They were mounted in a stereotaxic frame (Kopf, Tujunga, CA) and through a small craniotomy, they were injected into the right CA1 subfield (AP: 2.1mm, DV: 1.65 mm, ML: 2 mm) at a rate of 4 nl/s with 0.5 µl AAV1.Syn.GCaMP6f.WPRE.SV40 (titer: 4.65×I0_13_ GC/ml; Penn Vector Core). The injection pipette (Nanoject III, Drummond) was left in place for 5 min before it was slowly withdrawn. Thirty minutes later, a larger craniotomy was performed with a 1.8-mm diameter trephine drill and the overlying cortex was removed by suction. A gradient-index ‘GRIN’ lens (Edmund Optics, 1.8 mm diameter, 0.25 pitch, 670 nm, uncoated) was implanted, fixed with cyanoacrylate, and protected by Kwik-Sil (WPI, Sarasota, FL). The skull was covered with dental cement (Grip Cement, Dentsply, Long Island City, NY). The mice received one slow-release buprenorphine injection (s.c. 0.5 mg/kg), dexamethasone (s.c. 0.1 mg/kg) for 6 days, and amoxicillin in water gel for a week. Three to eight weeks later, once a good field of view was visible through the GRIN lens, the baseplate for a UCLA miniscope was implanted and fixed to the skull with dental cement (www.miniscope.org; Aharoni and Hoogland, 2019).

### Behavior

After recovery from surgery, animals were handled for a couple minutes 1-3 times a week until the baseplate was implanted. The animals were then habituated to wearing the miniscope in their home cage, first in the animal holding room and then in the experimental room. Once the animal was comfortable wearing the miniscope, we started the behavior experiment and recording.

The mice were exposed to two environments, a 32 cm-diameter circle with transparent walls and a 28.5 cm-side square with opaque black walls and distinctive orienting pattern on 3 of the walls. The surface area of the two enclosures were similar, within 2% of 800 cm^2^, but the floor of each environment was also distinctive; the circle’s floor was made of red plastic while the rectangle’s was of black metal. Orienting cues were also present in the room, and directly visible from the circular enclosure; the door and the animal transport cart were visible, and two salient cues were present on opposite walls of the room.

The mice explore each environment twice, for 5 min each time, in an interleaved fashion. The animal was placed in its home cage between trials for a couple minutes to allow us to change environments. Windows-based software (Tracker 2.36, Bio-Signal Group Corp., Acton, MA) determined the mouse’s location in every 30-Hz video frame from an overhead digital video camera.

This protocol was repeated for three consecutive days, every week of three consecutive weeks. For two animals, this was repeated in periods of 5 days (instead of 7). For all animals, we started calcium recording on the day the animal first explored the two environments. We aimed to record the entirety of the 5-min visits to the circle and square environments and 5-min sessions in the animal’s home cage were also recorded each day before and after the 4 sessions in the two experimental environments.

### Calcium recording and signal extraction

Neural activity data from 6 of the 41 mice met the quality analysis requirements described below and were analyzed to address the central question. When recording, the UCLA miniscope was attached to the baseplate, and fluorescent images from CA1 were recorded through the GRIN lens using the UCLA miniscope data acquisition hardware and software (www.miniscope.org). Thirty images were collected each second and analyzed offline. Before analysis, we screened the average calcium level of each recording and removed any day or week where large and abrupt changes to the mean fluorescence were observed. The images from each recording were aligned using the NoRMCorre algorithm (Pnevmatikakis and Giovannucci, 2017). The alignment was done separately for each recording but all recording alignments from a single mouse used the template generated by the alignment of the first video recorded that day. Aligned videos were then subsampled to 10 Hz by averaging and the recordings from each day were concatenated into a single file. Action potential activity was then estimated separately for each day from the ΔF/F GCaMP6f signal using the CNMF-E algorithm (Pnevmatikakis et al., 2016; Zhou et al., 2018), which simultaneously separates the cells’ fluorescence and deconvolves the calcium transients to infer spiking activity. Accordingly, we refer to ‘activity’ rather than ‘firing’, especially because calcium transients are likely to reflect bursts of action potentials rather than single action potentials (Ledochowitsch et al., 2019). The spatial footprint of the cells were seeded using a peak-to-noise ratio determined manually for each recording session. Units with high spatial overlap (>0.65) and high temporal correlation (>0.4) were merged and the CNMF-E algorithm was updated until no pair of units meet the criteria. Ensembles were evaluated manually for quality control to reduce the likelihood that artifacts were identified as cells. Furthermore, recordings were examined for evidence of photobleaching and minutes scale downward or upward trends in activity by regression analysis of the activity timeseries of individual cells for every one of the ≤54 (cylinder, box, home cage) recordings (6/day X 3 days x 3 weeks). Only recordings with no significant linear trends and without abrupt decreases in activity levels were studied.

To evaluate the integrity of the neuron ensembles we recorded, we used two metrics: temporal independence and spatial dependence. Temporal independence evaluated for each cell the average cell-pair correlation with all cells whose center is within 20-µm of the soma of the cell. Spatial dependance measured the proximity and overlap with nearby cells. For each cell, the CNMF-E algorithm computes a spatial footprint. The spatial dependence is the proportion of the summed footprints within a 20-µm radius area around the soma that is not attributable to the cell itself. These metrics confirmed to the best of our abilities, that no corrupted unit or ensembles were included in the dataset. Ensembles ranged in size from 39 to 588 isolated units, with half larger than 260 cells and less than 10% smaller than 100 cells.

### Alignment and cell matching

To match cells across days, every pair of recording days was first aligned to each other using the template used for the within-day alignment. This alignment was done using a non-rigid optical flow algorithm (Farnebäck, 2003), calcOpticalFlowFarneback function in the cv2 python library) whose parameters (window size, flow levels, number of iterations) were optimized for each animal. Overlapping spatial footprints were then matched conservatively (matches were performed using the Hungarian algorithm (Kuhn, 1955) with a maximum distance considered or cost of 0.7). Alignment and cell-matching algorithms were applied as implemented in the register_ROIs function of the CaImAn project (Giovannucci et al., 2019). Day-to-day alignments were verified by eye and for each alignment an f1-score was computed as the ratio of the number of cells aligned to the total number of cells. Alignments with an f1-score below 0.3 were discarded.

### Place Cell Identification

We computed cell-specific spatial activity maps from each cell’s activity, extracted at 10 Hz, and the animal’s location, subsampled to 10 Hz, by computing the average rate of the cell in each ∼2.5 x 2.5 cm bin. Linearized spatial activity maps were computed by changing the Cartesian coordinates to circular coordinates, with the origin at the center of the environment. We then computed the average rate of the cell in each 12° bin with the distance from the origin ignored.

To identify place cells, we used two metrics: Spatial coherence and information content (Muller and Kubie, 1989; Skaggs et al., 1993). Spatial coherence was defined for each cell as the correlation between the average rate at each location and the average of the average rate at the locations neighboring this location (up to 8). Information content was defined for each cell as:

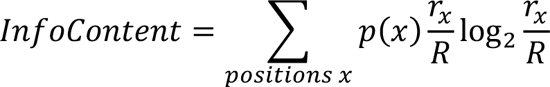

where *p(x)* is the probability of the animal being at postion x, *r_x_* the average rate at position *x* and *R* the average rate of the cell.

For each cell we computed both metrics as well as a distribution of randomized values with shuffled activity. The metric *val* was considered significant if

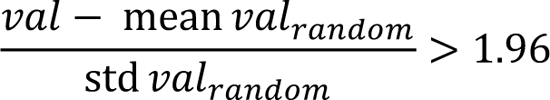

with *val_random_* being the distribution of randomized values.

We classified a cell as a place cell only when both metrics were significant.

### Place field similarity

The stability of a cell’s spatial tuning was assessed by comparing the spatial activity map in a pair of recordings. The Pearson correlation was computed between the pairs of activity rates at corresponding pixels in the activity rate maps. The average correlation (r) is reported for summary statistics but for statistical comparisons 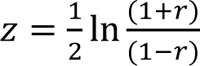 Fisher’s transform was used.

### Cell pair correlations

Kendall tau (τ) correlations were computed using 1-s time bins because this measure of association is robust to time series with a low range of values and many ties (e.g., 0 and 1). We evaluated cell pair correlations as a function of the distance between neurons and found that neurons within 25 µm of each other were, on average, more strongly correlated. Since our recording method does not allow us to determine whether this is physiological or an artefact, we decided to exclude all such pairs. Therefore, for all analyses using cell pair correlations, we excluded cells pairs that were not spatially separated (distance < 30 µm). We also excluded cells with very low activity (active less than 2% of 1-s time bins) as Kendall correlations computed on such time series have little meaning.

### Position-tuning independent (PTI) rate

From the rate maps (average rate at each position), we computed the expected rate as the average rate at the animal’s location / the sampling rate (here 100 ms). The expected rate is then binned into 1-s bins and subtracted from the 1-s binned observed rate.

### Support vector machine (SVM) decoding

SVM analysis was performed on concatenated 1-s binned activity time series from both environments using the sklearn python library. Training and decoding are performed 100 times on each dataset using different randomly separated training set (2/3 samples) and a testing set (1/3 samples). Decoding accuracy and features (cells or cell pairs) weights are then averaged across the 100 repetitions. This ensured that the results were not sensitive to random variations in the dataset split. Decoding accuracy is computed as the number of right decisions in the testing set. When decoding and comparing weights across day, no cross validation was used, and the entire reference day dataset was used to train the decoder. When decoding location, the SVM decoder was trained and evaluated using the 1-s activity vectors or coincidence vectors while the animal was in each of the two binned locations comprising the n(n-1)/2 pairs of bin locations, in each environment. The location decoding score was computed as the average of correct choices between the n(n-1)/2 pairs of locations. When decoding environment, the SVM decoder was trained and evaluated in the 1-s activity vectors or coincidence vectors during each day (two recordings in the cylinder environment and two recordings in the square environment). The environment decoding score was computed as the average of correct choices, using 100-fold cross-validation. Because SVM decoding is sensitive to the number of elements (i.e., vector dimension) for comparisons of different types of input, the input data were downsampled to the lowest dimension vector that was being compared.

### Network consistency

The Network consistency (van Dijk and Fenton, 2018) describes how well the momentary covariance in cell-pair activity fluctuations align to the overall activity correlation of the cell pair. Network consistency of an ensemble of time series is calculated as:

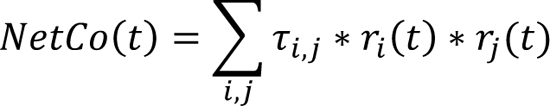

Where *τ_ij_* is the pairwise Kendall correlation between cell *i* and cell *j* and *ri(t)* the activity rate of cell *i* at time t. To compare network consistency values between animals we normalized it in each recording to the standard deviation of the distribution of values obtained when the identities of the cell pairs were shuffled 100 times. In the figure, we report the average network consistency for each recording.

### Correlation participation

To quantify the role of cell pair activity correlation in the SVM and IsoMap analyses, we evaluated for each cell their participation in correlations of different ranges of correlation t values. For each cell, we computed its participation in a range of t values as the number of times a cell participates in a pair with a t value within that range, divided by the total number of pairs in which the cell participates. In effect, it is the proportion of t values falling in a certain range amongst all cell pairs the cell participate in. In practice, this normalization has limited effect but allows us to remove any impact from the exclusion of some cell pairs.

### Geometric properties of high-dimensional neural manifolds

As detailed in (Chung et al., 2018b) neural manifolds are defined by D+1 coordinates: one to describe the location of the manifold center, and D axes that define the manifold variability. In this work, we consider that each of the P manifolds is sampled M times along N different features. In this work, P manifolds correspond to P recordings of the animal in each environment (square and/or circle), and N features correspond to N neurons.

To characterize the separability among neural manifolds, we introduce the concept of classification capacity, α_5_. Classification capacity is understood as the maximum number of manifolds that can be linearly separated given random assignment of binary labels to manifolds, and mathematically as the ratio between the critical number of manifolds, *P5* and the number of features, N. Borrowing the concept of support vectors from the problem of linearly separable points, in manifold theory, the high dimensional separating hyperplane is defined by a linear combination of what we call manifold anchor points. Each manifold contributes with (at most) a single anchor point, which uniquely define the separating plane between manifolds. The identity of the anchor points depends on the location and orientation of the manifolds as well as the randomly assigned binary labels. As a result, for a given manifold, there is a statistical distribution of anchor points. From the distribution of anchor points, we can introduce two geometric properties of manifolds: the effective radius, *R6*, defined as the square root of the total variance of the anchor points, normalized by the average norms of the manifold centroids, and the effective dimension, *D6* which captures the dimensionality of the anchor points along the different manifold axes. We define the manifold total extent, a measure of the space occupied by the high dimensional neural manifold as the mean *R6JD6*.

According to the replica mean field theory of manifolds, larger *R6*, *D6* and *R6 JD6* of the manifolds are linked to lower linear separability between these manifolds, indicating more overlap between them.

### Betti Numbers and dimensionality reduction

Principal Component Analysis (PCA) and IsoMap dimensionality reduction were performed each day separately on concatenated 1-s binned activity time series from both environments as well as the home cage, filtered with a Gaussian filter of 1-s standard deviation. Both transformations used the sklearn python library. IsoMap was performed using Euclidian distance and 5 neighbors (Balasubramanian and Schwartz, 2002; Tenenbaum et al., 2000). Betti barcodes were computed each day on a 10-dimension IsoMap projection using the Ripser algorithm (Tralie et al., 2018) in the scikit python library. To compute isometries of Betti number 2, the data were randomly subsampled to 700 data points (∼40%) to allow for computational tractability.

### Low-dimensional neural manifold analysis

We start by considering a N-dimensional neural manifold. The dimensionality of the neural manifold is evaluated using the participation ratio (PR), defined as:

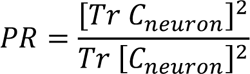

where *Tr C_neuron_* is the trace of the covariance matrix of a pattern of firing rates across M recorded neurons at some specified time window (Gao et al., 2017).

To quantify the anti-cofiring properties of each cell, we introduce the anti-cofiring power, defined as the number of cell pair correlations with τ < −0.05, normalized by the total number of correlations calculated for the cell (i.e. n-1).

Further, to describe the distribution of distances to the centroid in a N-dimensional manifold, we employ two different measures. First, we use the skewness as a measure of the asymmetry of the data around the sample mean, i.e., a measure of the tail-ness of the distribution, which contains the ‘outliers’, earlier shown to be most anti-cofiring cells, defined as:

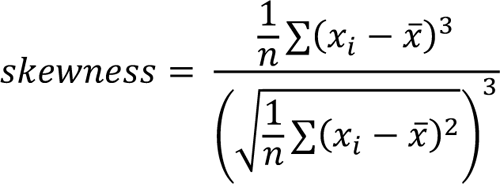

Second, to measure the similarity between distributions for different ensembles, we use Kullback-Liebler divergence (KLD). KLD is a measure of how one probability distribution f_0_ is different from a second, reference probability distribution f_2_. This measure can be interpreted as the expected excess surprise from using *f_0_* as a model when the actual distribution is f_2_.

Then, we construct 2D neural manifolds using IsoMap projected neural activity into a three-dimensional subspace. The 2D manifolds are constructed using the in-built function *alphaShape* in MATLAB. The algorithm creates a 3D bounding volume using a triangulation algorithm. To identify the overlapping coefficient between two environments, we first identify the number of points from one environment manifold that belongs to the other environment manifold, using the in-built command *inShape*. We then compute the overlapping coefficient (overlap) by normalizing the number of occurrences from Environment 1 inside the 2D manifold of Environment 2, with the total number of points from Environment 1.

To explore the multi-stable planar manifold hypothesis, we compute the best fit plane to the 3D IsoMap projected point cloud data for each environment using an in-house MATLAB function. The algorithm fit 1-s activity vectors in a 1 min window that was advanced in 1-s increments. The algorithm fits a least square solution for the normal vector that defines the best fit plane. The algorithm minimizes the sum of the squared dot product of a trial normal vector with a vector passing through the centroid of the data. The output is a normal vector and the centroid of the 3D point cloud data.

To further quantify the distinction between environments, we compare the distribution of angles measured between the normal vectors of each pair of consecutive planar fits within a particular environment. These distributions were compared between repeated recordings of the same and different environments using Kuiper’s statistic, a rotation-invariant Kolmogorov-type test statistic.

Finally, to describe the trajectories inside the neural manifold, we introduce two quantities: localness, defined as the Euclidean distance between two temporally adjacent points, and smoothness, defined as the angle between the normal vectors of two temporally adjacent planar fits.

### Statistics and visualization

All data were plotted using boxplots unless the distribution contains less than 10 data points, in which case individual data points are plotted. For all boxplots, the height is determined by the interquartile range and the median is indicated with a line. The whiskers extend to the furthest data point up to 1.5 times the interquartile range past the boxplot limit. Data points past the whisker limits are either plotted individually or omitted when too large for better visual representation.

Statistically significant differences are indicated on the plots. One asterisk indicates p < 0.05 (or the corresponding corrected value) and two asterisks indicates p < 0.01 (or corresponding corrected value). When indicated, a statistically significant effect of the factor week is indicated as a line above the graph with asterisk(s).

Within subject comparisons were performed, when possible, using paired statistics such as Student’s paired t test, and the corresponding non-parametric test, Wilcoxon test. Comparisons amongst multiple factors or amongst more than two groups were performed using ANOVA, with repeated measures (RM) when the effect of week was being evaluated. The Hotelling-Lawley correction was used when the data being compared by RM ANOVA violated the sphericity assumption as assessed by Mauchly’s Test of Sphericity. For analyses in which a particular percentage subset of the data was included or excluded, the factor of percentage was treated as a continuous variable with df = 1. Dunnett’s tests were used to evaluate post-hoc pairwise comparisons against a control, such as an initial week or control subset, when appropriate. Bonferroni-corrected multiple comparisons were used when only a subset of pairwise comparisons were of interest and when comparing multiple groups against a value or control group (if not present in the initial RM ANOVA). For comparison amongst multiple factors or amongst more than two groups where the data was not normally-distributed we performed Friedman’s statistical test and Kruskal Wallis test. All correlation data were Fisher-z transformed before statistical analysis using parametric methods. The test statistics and degrees of freedom are presented along with the p values and effect sizes for all comparisons, with only power of ten indicated when a value is below 0.001. A value of p < 0.05 was considered significant. The statistical analyses of data in a figure are presented in the corresponding figure legend.

### Large Environment model

Larger place field maps were created from the concatenations of the place field map of four different cells. Only cells that were categorized as place cells in at least one repetition of the two environments were considered. For each larger map, we obtained four firing field maps, one for each visit of the two environments. Cells from a single day (day 6) and from all six animals were used to create the 151-‘composite’ cell ensemble. Results did not significantly change across days.

A random walk was then simulated across the larger environment but limited to different boundaries depending on the environment size evaluated. To simulate the random walk, we started at a random position within the allowed boundaries and randomly sampled a move every dt=0.1s. The size of the large place field space was considered to be 24×24 space units. Position change was sampled randomly from [-1, 0, 1] and multiplied by a speed variable. The speed variable was initiated at 0.1 and, similarly to the position, was changed at each time step by a random sample from [-0.1, 0, 0.1], with 0.1 and 1 being the speed upper and lower bounds respectively. This corresponds to a maximum speed of 2.5 to 25 cm/s in real space. Four different random walks were created for the two visits to the two environments and expected activity was created for each place map as described in the *Position-tuning independent (PTI) rate* method section. Environments were then decoded as described in the *Support vector machine (SVM) decoding* method section.

### Spike-time dependent plasticity network model and analyses

The code implementing the spiking neural network model was custom written and has been made freely available (https://github.com/william-redman/EI-STDP-place-code-network).

### Architecture

As illustrated in Fig. 1B, the spiking neural network was organized as follows: The input to the network came from a layer of n_input_ = 1000 neurons that were tuned to locations on the track. This population fed into a layer of n_excit_ = 500 excitatory neurons with weights W^InputE^. The connection between the input and excitatory populations were non-plastic. The excitatory population was both recurrently connected to itself (W^EE^), and to an inhibitory population of n_inhib_ = 50 neurons (W^EI^). This inhibitory population is connected back to the excitatory layer (W^IE^). The E→E, E→I, and I→E weights were all plastic.

At initialization, all values of W^InputE^ were uniformly sampled from the interval (0, *W^InputE^_max_*). For W_EE_, each weight either took a uniformly sampled value from the interval (0, *W^EE^_max_*), with probability 1 – p_EE_, or was set to 0, with probability p_EE_. The diagonal entries of W^EE^ were all set and kept at 0. To ensure that the total amount of initial synaptic weight was conserved across the different weight populations, we set *W^EE^_max_* = *W/^InputE^_max_/p_EE_*. The same initialization procedure was repeated for W^EI^ and W^IE^ with *W_max_^EI^ = W_max_^InputE^/p_EI_ and W_max_^IE^=W_max_^InputE^/p_IE_*.

### Input layer

The input layer to the spiking neural network was made up of n_input_ neurons, each tuned to at least one position on the linear track. The distribution of number of locations each input neuron was tuned to was binomial, with p_input_ = 0.20. The centers of these tunings took a continuous value from [0, 10). The tuning to each center was taken to be Gaussian. That is, for neuron *i* with center, *c_t_*, the tuning takes the form

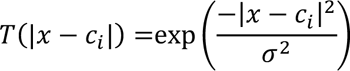

where |·| is circular distance and *x* is a given location on the track. Neurons with n > 1 center take the sum of their tuning, which then takes the form

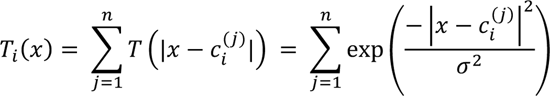

The probability of input neuron *i* firing when at position **x** is proportional to the input with added noise such that, at any given time point, it is given by

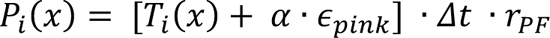

where *Δt* is the time step length of the simulation, *r_PF_* is the “in-field” firing rate, and *e_pink_* is a noise term that is drawn from a pink noise distribution (https://github.com/cortex-lab/MATLAB-tools/blob/master/pinknoise.m) and is weighted by *α.* In the case of a negative noise term that is larger than *T!* (x), *P!* (x) was set to 0.

### Neuron model

The neurons in the excitatory and inhibitory populations were modeled as simplified integrate-and-fire neurons. That is, the voltage of excitatory neuron ¿ at time *t* is given by the weighted sum

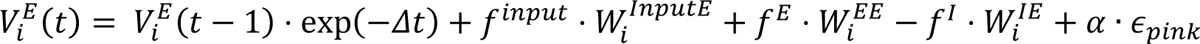

where *f^mput^* is the spikes of the input population and *w¦^nputE^* are the weights from the input layer to neuron ¿ (and similarly for E and I being the excitatory and inhibitory populations, respectively). As in the case of the input layer, *e_pnık_* is a noise term that is drawn from a pink noise distribution, which is weighted by α. When the voltage is greater than the threshold *T_ļF_*, the voltage is set to zero and *fE* is set to 1 for the next time step. We include an absolute refractory period, *τ_IF_*, where *V_E_* is kept at 0.

For inhibitory neuron *i*, the voltage is given by

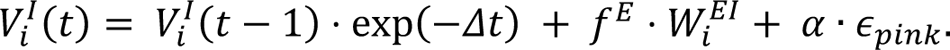

### STDP Plasticity

All E→E, E→I, and I→E weights could be modified via spike-timing dependent plasticity (STDP) rules. In the experiments where one (or all) of the plasticity types were removed, then those weights were not subject to STDP, but were instead frozen.

The recurrent excitatory weights were modified at each time step via excitatory STDP that has the following form

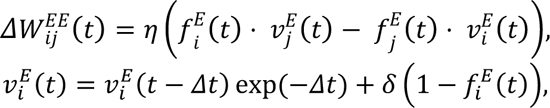

where the convention that *i* is the post-synaptic neuron and *j* is the pre-synaptic neuron is followed. ſ*_i_^E^(t)* is the spiking activity of the excitatory neuron *i* at time *t*, η is the learning rate (and maximum amount of change allowed at each time step), and δ is the Kronecker delta function. The second equation keeps track of the spiking history of each neuron.

The weights from the excitatory population to the inhibitory population were similarly modified

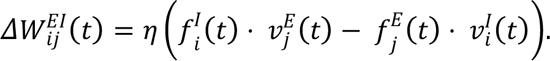

Finally, the weights that projected from the inhibitory population to the excitatory population were modified at each time step via inhibitory STDP that has the following form

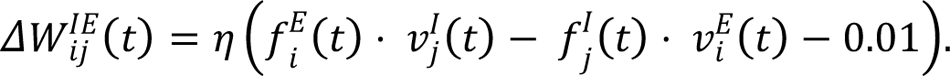

As negative weights do not make physical sense in our implementation of the spiking neural network, at each time step we enforced the condition (W < 0) = 0 for all weights. Similarly, we enforced that (W > *W_max_*) = *W_max_*.

### Single track experiment metrics

To evaluate the ability of a network with a given set of plasticity conditions, we ran 10 networks, each with a different set of random initial weights and different tuning centers for the input neurons, for 100 laps. To get a qualitative understanding, we split the track into 10 discrete locations and computed the spatial activity rate maps from the experiments using the last 25 laps (examples shown in Fig. 1C top). To make the observations we derived from the rate maps quantitative, we also computed two different similarity metrics for each excitatory neuron every 10 laps (using *only* the activity from those 10 laps). Both have the form

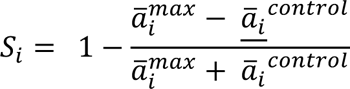

where *<u>a</u>^max^_i_* is the average rate at the location with maximal activity and *<u>a</u>^contro1^* is the average rate for a control location. For nearest neighbor similarity, the control locations are the positions on either side of the maximal location (circularly for boundary locations). For halfway similarity, we take the position on the track that is (circularly) the farthest from the location of maximal activity. The mean and standard error of the mean (SEM) for each of these metrics are plotted in Fig. 1C bottom.

### Two track experiments

To investigate how the spiking neural network was affected by exposure to a novel environment, and to see what role plasticity played in the encoding of the novel environment, we initialized and ran 10 networks under different plasticity conditions. For each experiment, we created two tracks, track A and track B, defined by their random input neuron tunings (i.e., T_i_), with the weights that connected the input to excitatory neurons (i.e., W^InputE^) kept constant for both tracks. We trained a naïve network on track A for 30 laps (the number of laps needed for the similarity metrics to reach a plateau), then transferred that network (that is, used the “learned” W^EE^, W^EI^, and W^IE^) to either track A again, called A’, or to track B to compare the activity obtained from the re-exposure to the “same” environment, track A, to the exposure to a “different” environment, track B. This investigated how much of the difference between the activity on track A and track B was due to noise and the stochastic nature of the spiking network.

To examine the effects of plasticity and novelty quantitatively, we computed two metrics. The first was the Pearson correlation of the rate maps computed on the last 50% (15 laps) of the activity (Fig. 1A left). We call this place field similarity, and the mean and SEM of the 10 simulations are plotted in Fig. 1D, top, for each plasticity condition. The second metric is the Pearson correlation of the Kendall correlations computed on each excitatory neuron cell pairs, again computed on the last 50% of the activity (15 laps) (Fig. 1A right). We call this cofiring similarity, and the mean and SEM of the 10 simulations are plotted in Fig. 1D, bottom, for each plasticity condition. To compute Kendall correlations, we first split the activity into bins of length *Δt_Kendalı_ =* 0.25 and then averaged within each bin.

### Parameters

The parameters used for the simulations are listed below. For more details on the implementation of the spiking network, see the freely available commented code (https://github.com/william-redman/EI-STDP-place-code-network).

| Parameter | Value | Description |
| --- | --- | --- |
| $n_{input}$ | 1000 | # input neurons |
| $n_{excit}$ | 500 | # excitatory neurons |
| $n_{inhib}$ | 50 | # inhibitory neurons |
| $p_{input}$ | 0.20 | parameter of binomial distribution for number of tuning centers of input neuron |
| $p_{inputE}$ | 1.0 | parameter of binomial distribution for number of excitatory neurons each input neuron is connected to |
| $p_{EE}$ | 0.25 | parameter of binomial distribution for number of excitatory neurons each excitatory neuron is connected to |
| $p_{EI}$ | 0.175 | parameter of binomial distribution for number of inhibitory neurons each excitatory neuron is connected to |
| $p_{IE}$ | 0.30 | parameter of binomial distribution for number of excitatory neurons each inhibitory neuron is connected to |
| $\eta$ | 0.0005 | synaptic learning rate |
| $\alpha$ | 0.05 | strength of pink noise |
| $\Delta t$ | 0.007 | time scale of simulation |
| $r_{PF}$ | 8 | firing rate at tuning center |
| $\sigma^2$ | 0.1 | width of tuning center |
| $\tau_{IF}$ | 0.02 | absolute refractory period of excitatory and inhibitory neurons |
| $T_{IF}$ | 1 | spiking voltage threshold |
| $\Delta t_{Kendall}$ | 0.25 | size of bin used to average the activity before Kendall correlation |
| $W_{max}^{InputE}$ | 0.05 | maximum initial weight between input neurons and excitatory neurons |
| $W_{max}$ | 0.50 | maximum value any weight can take |

### Histology

Mice were deeply anesthetized with Nembutal (100 mg/kg, i.p.) and perfused through the heart with cold saline followed by cold 4% paraformaldehyde (PFA) in PBS. The brains were removed, postfixed overnight in 4% PFA, and then cryoprotected in 30% sucrose for a minimum of 3 days. The brains were sectioned at 30–50 µm, stained with DAPI, and examined with a fluorescence microscope.

