## Supplemental Information for "A manifold neural population code for space in hippocampal coactivity dynamics independent of place fields"

[=] Equal Contribution

### Supplementary figures

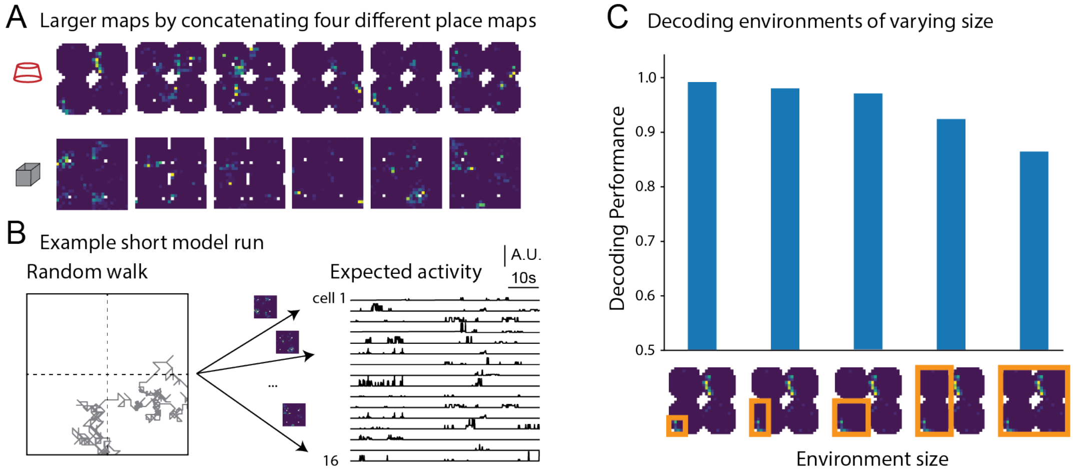


**Figure S1. Evaluation of the impact of multiple fields on decoding environments from place cells.** (A) Example of corresponding larger maps in two environments. Each map is created by concatenating the place maps of four place cells in the circle and square environment respectively. (B, left) Example of the random walk in the large environment during a simulation corresponding to 1 minute and (B, right) example 16 'composite' cells from a 152-'composite' cell ensemble. From each map, an expected trace is created from the random walk position and the average rate at the corresponding position. (C) Performance for decoding two repetitions of each of the two environments depending on the environment size in a 240-minute-long simulation. Decoding performance was stable for simulations of 60 to 240 minutes.

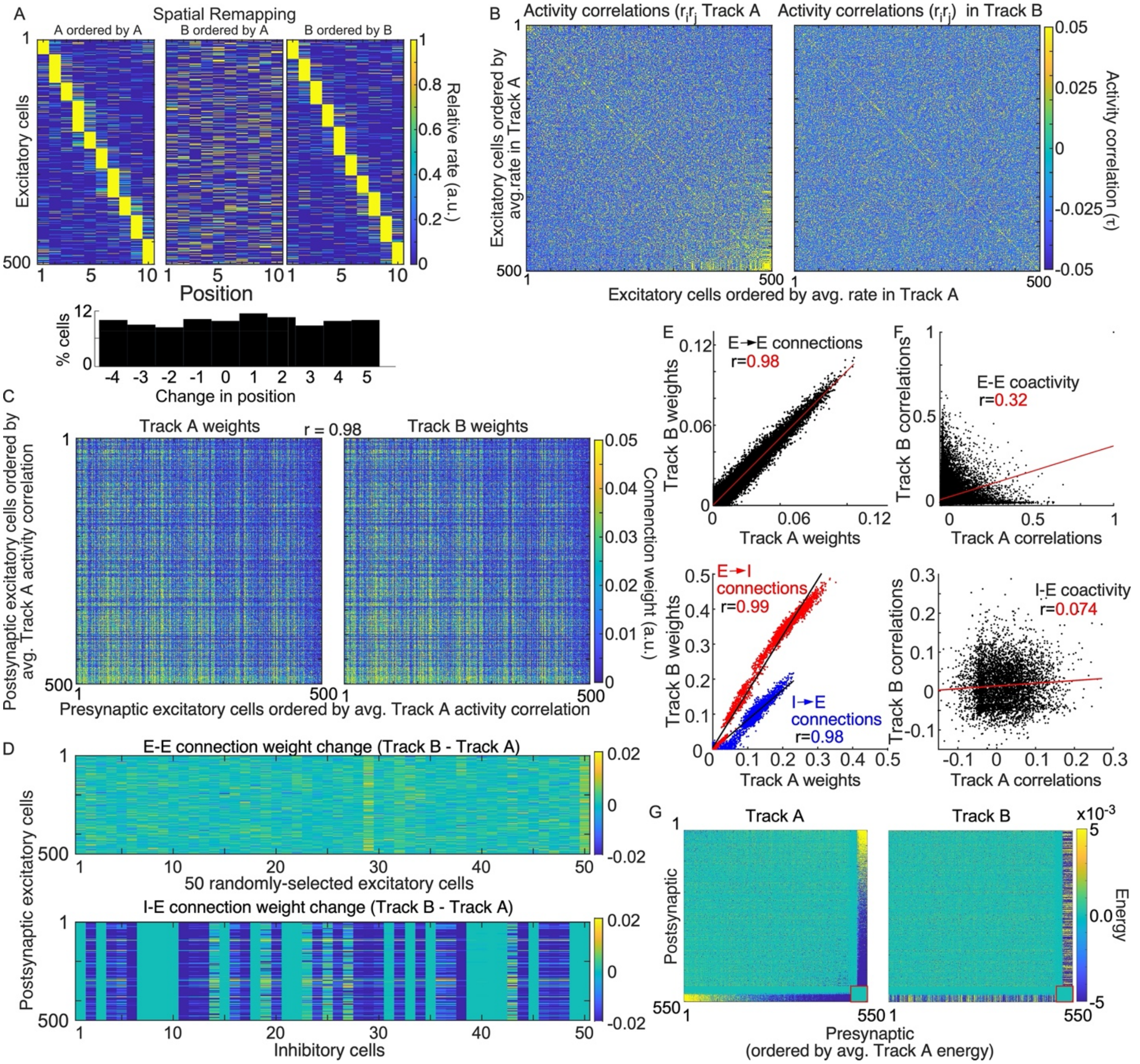


**Figure S2. Related to Figure S1. Network model predictions that reregistration underlies place field remapping.** (A) The STDP rules generate cell-specific location-specificity with elevated firing tuned to single "place field" locations in a small environment that reorganize to random locations (bottom) when the environmental inputs are remapped. Compare location-specificity organized by activity on track A (left, middle) and Track B (right) locations of peak firing. (B) The corresponding remapping-induced changes of E-E coactivity ($r_i r_j$) and (C) E→E connection strengths ($w_i w_j$). (D top) The remapping-associated change of E→E connection strengths depicted for 50 randomly-selected E-E pairs illustrate the changes are modest and fluctuate around 0, whereas (D bottom) the changes are greater at I→E connections. Notice how each I unit develops a cell-specific global pattern of network connections. I Despite remapping and engaging the STDP rules, the E→E connection strengths (top) tend to remain unchanged, whereas (bottom) the inhibitory weights reorganize by I→E weights decreasing and the E→I weights increasing. (F) The result is E-E (top) and E-I coactivity reorganize. (G) Energy, computed for each E-E, E-I, and I-I cell pair in the two tracks. I→I energy is highlighted by a red box. E cell IDs are 1-500, I cell IDs are 501-550.

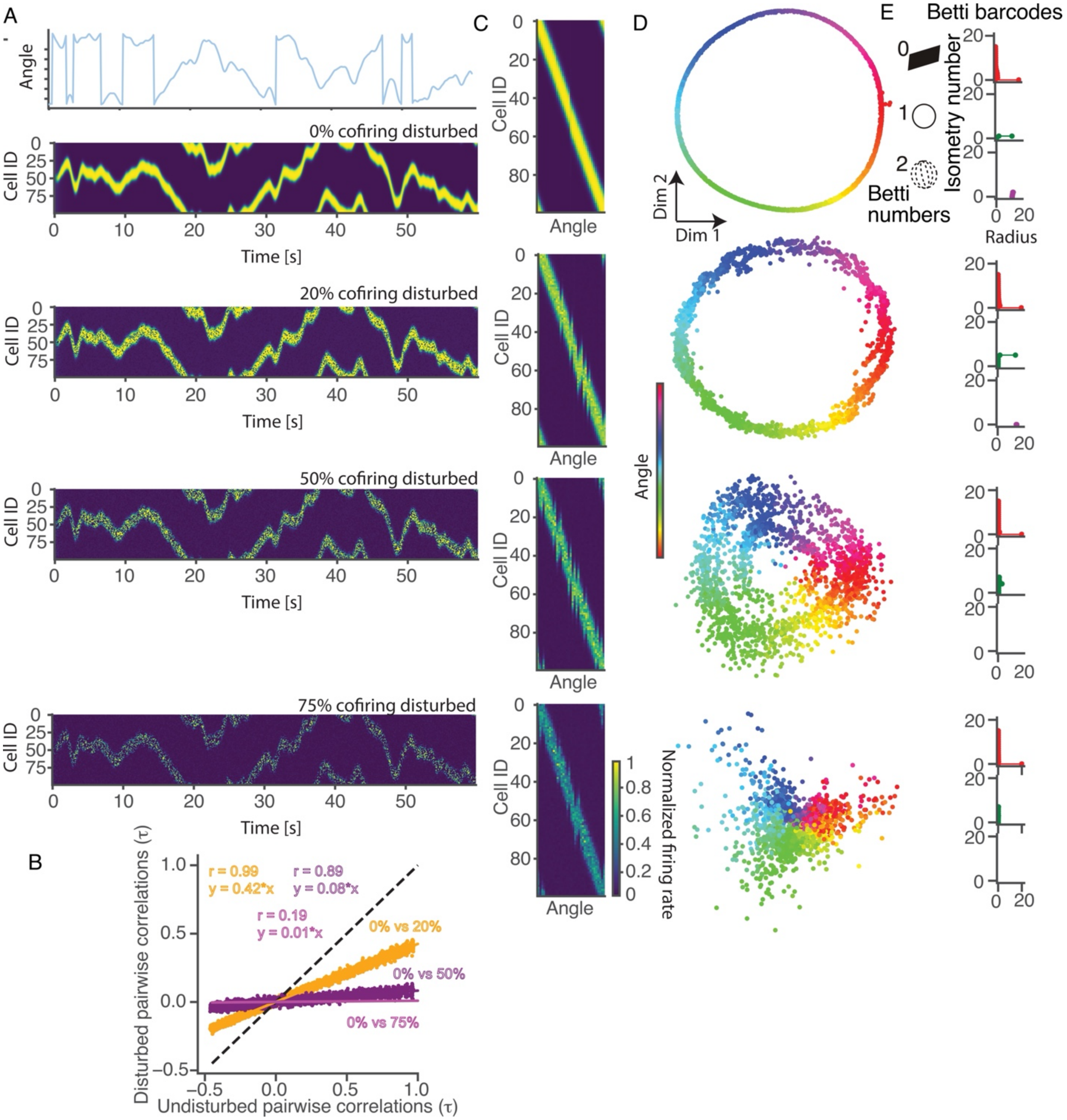


**Fig. S3. Related to Figure 3. Cofiring relationships determine population geometry of ensemble discharge.** Neural network simulation of spatial tuning along a ring environment illustrates that cofiring relationships determine population geometry. A) Normalized firing rates of 100 spatially tuned units during spatial exploration (top) with different levels of network disturbance. Cofiring during the indicated percentage of simulation moments was disrupted by randomly selecting an active unit and setting its activity to 0. B) Comparisons of all pairwise correlations confirming that the disruption effectively prevents reproducing the cofiring relationships of the undisturbed network. C) The disturbed cofiring does not corrupt spatial firing, D) but it does corrupt the ring structure of the population geometry as seen when projected in a 2-D IsoMap subspace.

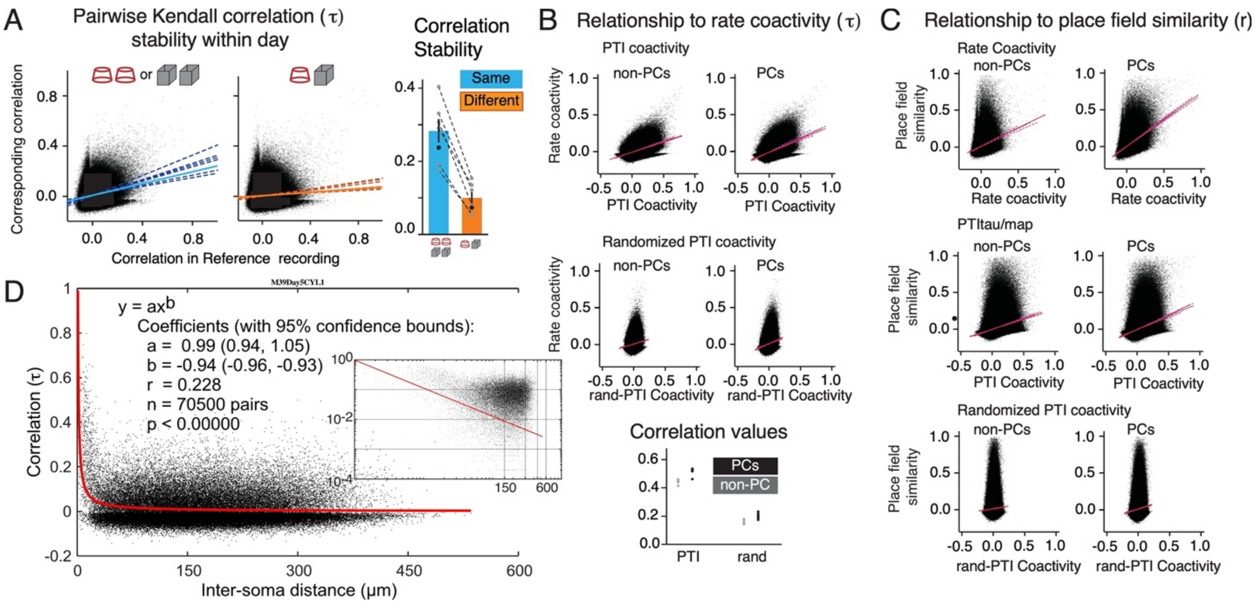


**Figure S4. Related to Figure 3. Additional Rate and PTI Coactivity information**
(A, top) Scatter plot of coactivity (τ) stability in (left) the same environments and (right) different environments. Dashed lines are regression lines for individual animals with (bottom) Pearson correlation (r) values plotted in grey over the bar graph indicating the average relationship. Solid lines indicate overall regression, with r values in black in bar graph. (B) Relationship between rate coactivity (τ) and (top) PTI coactivity or (middle) randomized PTI coactivity in scatter plots. Dashed mauve lines indicate regression for individual animals (bottom) with r values plotted in black for place cells and grey for non-place cells. Solid red lines indicate overall regression. (C) Relationship between place field similarity (r) and (top) rate coactivity, (middle) PTI coactivity, or (bottom) randomized PTI coactivity in scatter plots. Dashed mauve lines indicate regression for individual animals while solid red lines indicate overall regression. (D) Cell pair coactivity within a 376-unit ensemble recording during exploration of the cylinder on day 5, measured by correlations as a function of distance between the soma pairs. The data are fit significantly by a power law (inset: log-log plot).

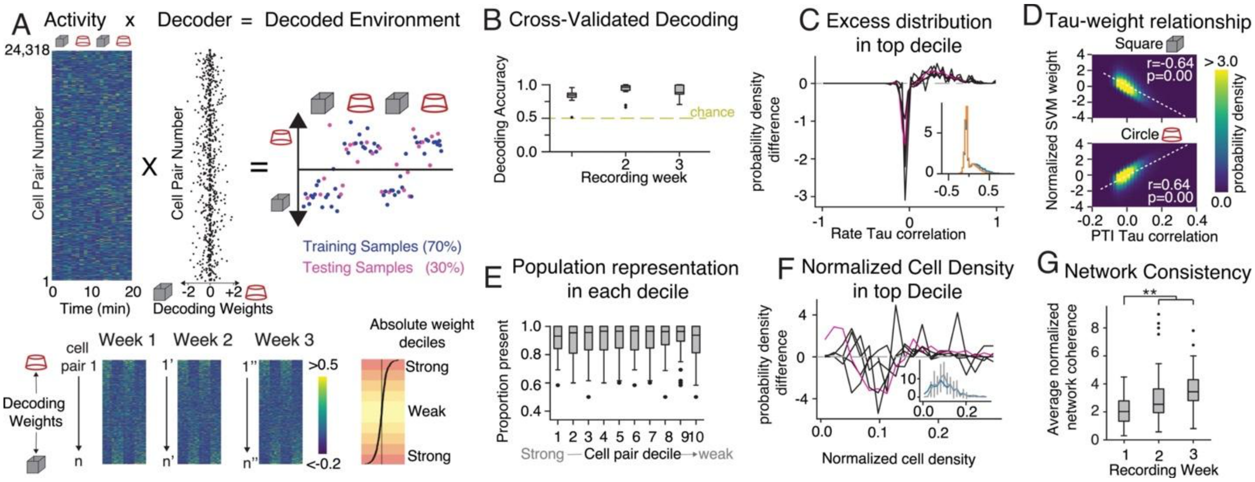

**Figure S5. Related to Figure 4. Decoding environments from rate coactivity.** (A) Illustration of SVM decoding from rate coactivity; (top) Rate coactivity, computed every 60 s, is projected with SVM weights onto a single dimension that separates the two environments. (bottom) Cell-pair coactivity from an example ensemble ordered by SVM weight each week; The absolute values of the weights, indicative of separation strength, are classified by decile. (B) Cross-validated decoding accuracy is very high for all ensembles. C) Difference between overall and first decile distribution of cell pair correlations (example distributions in inset and shown in purple in main figure) indicates importance of coactive cell pairs, confirmed by (D) 2-D distribution of SVM weights against Rate coactivity. (E) Proportion of individual cells present in at least one cell pair in each SVM-ordered cell pair decile. (F) Difference between random and first decile normalized distribution of individual cell participation in decile subset (example distributions in inset and shown in purple in main figure). (G) Rate network consistency normalized by the standard deviation of the cell-pair randomized distribution (Week: $F_{2,60}$ = 28.49, p=$10^{-9}$, $\eta^2$ = 0.13; Dunnett's tests: week 0 < weeks 2, 3).

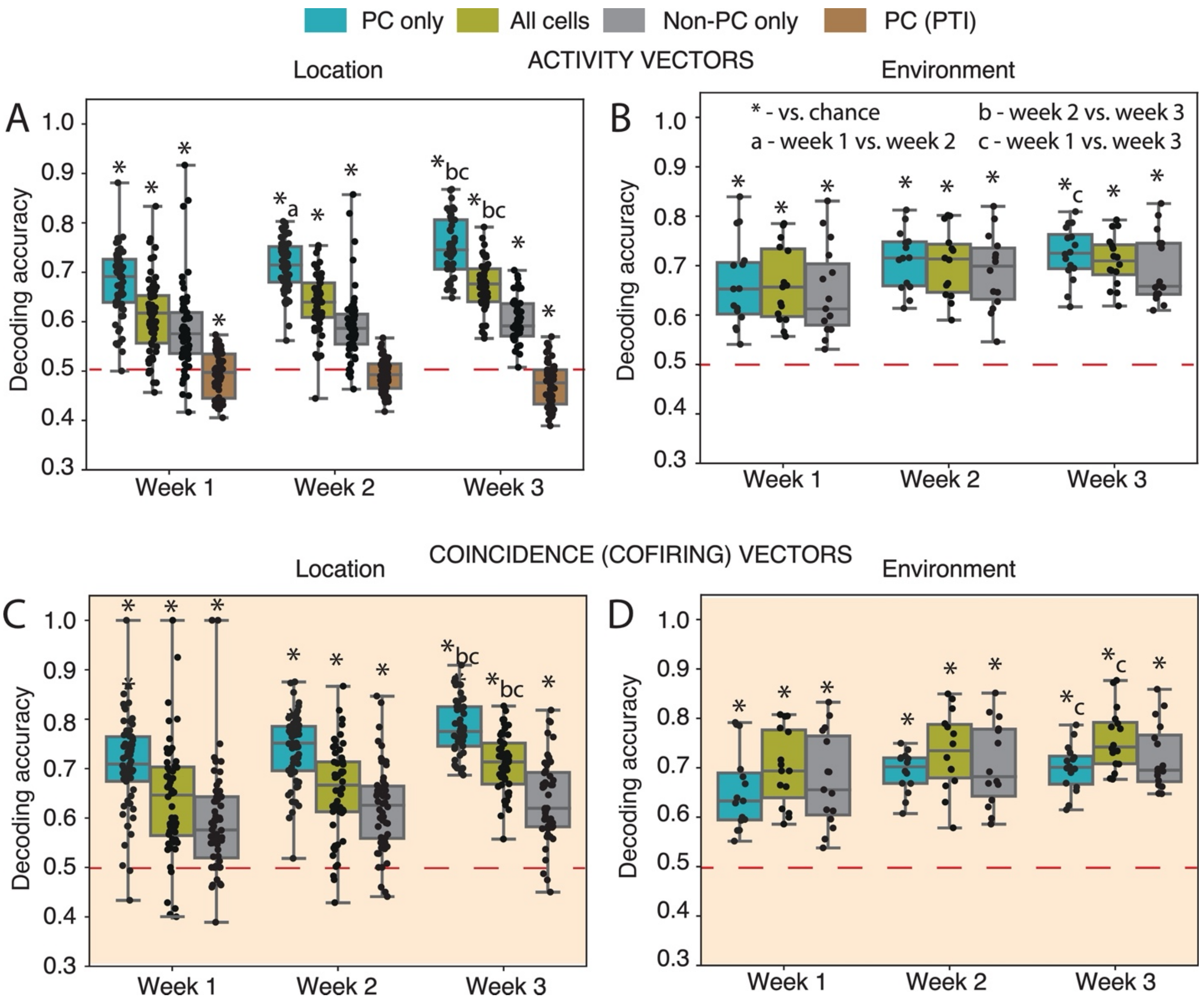


**Fig. S6 (related to Fig. 4). SVM-decoding of current location and environment from ensemble activity vectors and coincidence vectors constructed from functional subgroups of cells recorded across weeks.** Cells were classified as place cells (PC) and non-place cells (Non-PC), and the place tuning was eliminated from the place cells by computing position-tuning independent (PTI) timeseries. The ability to decode location from 1-s neural activity vectors was compared using data samples from the four cell classes: All cells, PCs, Non-PCs, and the PTI data sets. Because SVM decoding improves with number of features (cells), we subsampled to build activity vectors with the same number of cells as the PC group. A) Decoding location from 1-s activity vectors shows the effect of cell class is significant ($F_{2.52,\ 153.40} = 448.80$, $p = 10^{-71}$) as was the effect of week ($F_{1.74,\ 106.20} = 12.93$, $p = 10^{-5}$) and the interaction ($F_{4.57,\ 215.50} = 7.42$, $p = 10^{-6}$). Post-hoc Tukey's tests confirmed decoding accuracy with all the cells improved across weeks (week 1 = week 2 < week 3). This was largely due to the place cells, decoding from which also improved across weeks (week 1 < week 2 < week 3), whereas decoding from non-place cells did not change across weeks (week 1 = week 2 = week 3). Similarly, decoding from place cells after removing the place tuning (PTI) also did not change across weeks (week 1 = week 2 = week 3). PTI decoding accuracy was below chance (mean ± SD: 0.486 ± 0.043, comparing 0.5 to all outcomes collapsed across weeks; $t_{165}=4.224$, $p=3.96 \times 10^{-5}$). Decoding accuracy was better with place cells than all the cells when the number of cells used to train the SVM and decode were matched, which is why decoding with non-place cells was worse than with place cells, but higher than chance estimated by the PTI decoding ($t_{164} = 16.53$, $p = 10^{-37}$), as previously reported (Stefanini et al, 2018). B) We then investigated decoding the environment from 1-s activity vectors. The effect of cell class was significant ($F_{1.5, 22.49} = 11.37$, $p = 10^{-4}$), as was the effect of week ($F_{1.896, 28.44} = 3.65$, $p = 0.04$), but not the interaction ($F_{3.076, 39.22} = 0.48$, $p = 0.7$). Tukey tests do not distinguish the cell classes except in week 3 when decoding from place cells was better than from non-place cells. C) Decoding locations from cofiring relationships. The natural analysis is to decode location from the vector of cell pairs correlations like in Figures 4 and 5. However, estimating correlations on the timescale of seconds is not technically possible. Sampling for approximately one minute is necessary to estimate a cell pair firing correlation, whereas the animal changes location on the timescale of seconds. Nonetheless to investigate the ability to decode location from cofiring, we developed a 1-s time-scale coincidence-based analysis. Every one second whether a cell pair fired together (coincidence = 2), or one member of cell pair fired (coincidence = 1) or none of the cells fired (coincidence = 0) was computed for each cell pair, and the vectors of these coincidences were used to train and decode current location. Panel C shows the decoding accuracy from coincidence vectors assembled using the three cell classes (all cells, PC, non-PC). The effect of cell class was significant ($F_{1.72, 103,20} = 77.58$, $p = 10^{-19}$) as was the effect of weeks ($F_{1.89, 113.30} = 20.33$, $p = 10^{-8}$) but not the interaction ($F_{3.15, 145.11}= 0.80$, $p = 0.49$). Tukey tests indicate decoding by coincidence vectors from pairs of PC was better than coincidence vectors from the other cell classifications (PC > all cells > non-PC, in all weeks except week 1 and week 2: PC > all cells = non-PC). Decoding from coincidence vectors of non-PC and all cell pairs was higher than chance ($t_{163}$'s $\geq 15.38$, p's $\leq 10^{-33}$). Decoding from coincidence vectors of PC pairs and all cell pairs improved across the weeks (week 1 = week 2 < week 3) indicating that location decoding is driven and acquired by place cells. D) Decode the environment from 1-s coincidence vectors from pairs of all cells, PC, non-PC. The effect of cell class was significant ($F_{1.53, 22.95} = 25.32$, $p = 10^{-6}$), as was the effect of week ($F_{1.97, 29.55} = 3.58$, $p = 0.04$), but not the interaction ($F_{2.31, 29.47} =0.33$, $p = 0.75$). Tukey tests indicate decoding the environment from coincidence vectors is best when computed from all cell pairs across the weeks (PC < non-PC < all cells, except week 2: non-PC = PC < all cells). The decoding accuracy improvement across weeks is driven by place cells because decoding accuracy from non-place cell pairs do not change across weeks, but decoding accuracy improves across weeks from place cell pairs and all cell pairs (week 1 < week 3).

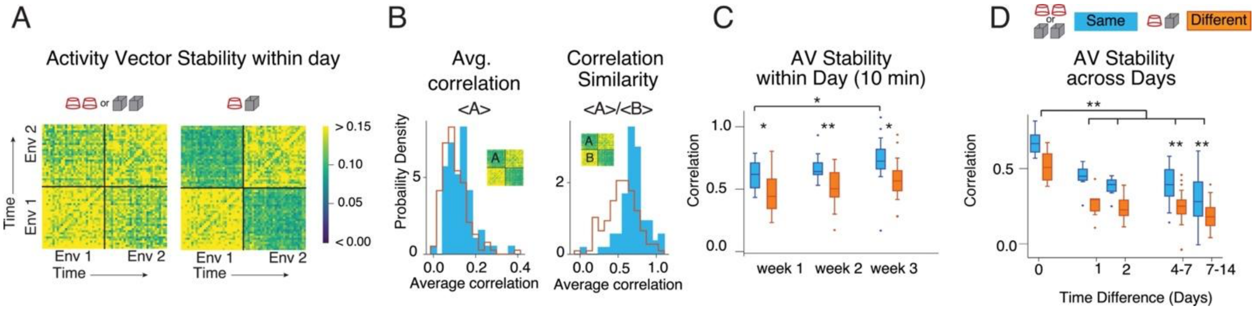


**Figure S7. Related to Figure 5. Activity vector stability and environment discrimination**
A) Illustration of activity vector correlation matrices, comparing similar or distinct environments, where each pixel is color-coded by the correlation between the 2 activity vectors observed at the time indicated on the x- and y- axes. (B) Histogram indicating distribution of (left) average correlation between activity vectors of separate recordings and (right) 'correlation similarity', average normalized by average correlation between activity vectors of the same recording. (C) Within-day correlation similarity improves across weeks (Week: $F_{2,28} = 4.47$, $p = 0.021$, $\eta^2 = 0.081$; Env. Change: $F_{1,29} = 24.10$, $p = 10^{-5}$, $\eta^2=0.20$; No interaction: $F_{2,28} = 0.18$, $p = 0.83$, $\eta^2 = 10^{-4}$; Dunnett's tests: week 1 < weeks 3 for similar environments; Same vs. Different is significant each week, *p's<0.015, **p = 0.0017). (D) Across-day correlation similarity degrades over time (Time: $F_{4,132} = 26.28$, $p = 10^{-16}$, $\eta^2 = 0.37$; Env. Change: $F_{1,132} = 32.99$, $p = 10^{-8}$, $\eta^2 =$

0.15; Interaction: $F_{4,132} = 0.40$, $p = 0.81$, $\eta^2 = 0.043$; Dunnett's tests vs day 0 **p<0.007; Same vs. Different **p ≤ 0.0013).

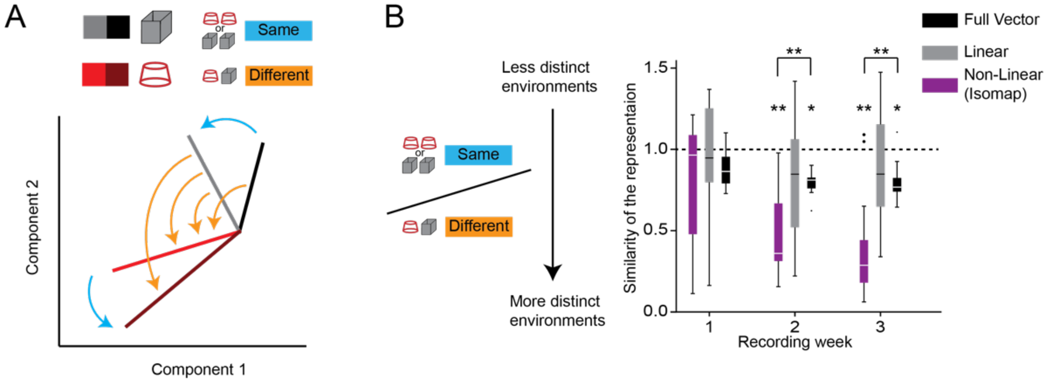


**Figure S8. Related to Figure 7. Computing the similarity of representation manifolds.** (A) Illustration of average 2-D vectors computed from 1-s individual activity vectors, recorded during the exploration of the two environments, projected on a 2-D subspace. Repetitions of the same environments are shown as different shades of the same color. The vector difference is calculated between the resultant 2-D vectors from recordings in the same environment (blue arrows) and from recordings in distinct environments (orange arrows). (B) The ratio of the same-environment vector difference over the different-environments vector difference measures the similarity of the representations shown in Fig. 6D (reproduced here), where smaller values indicate that the different environments are more distinctly represented

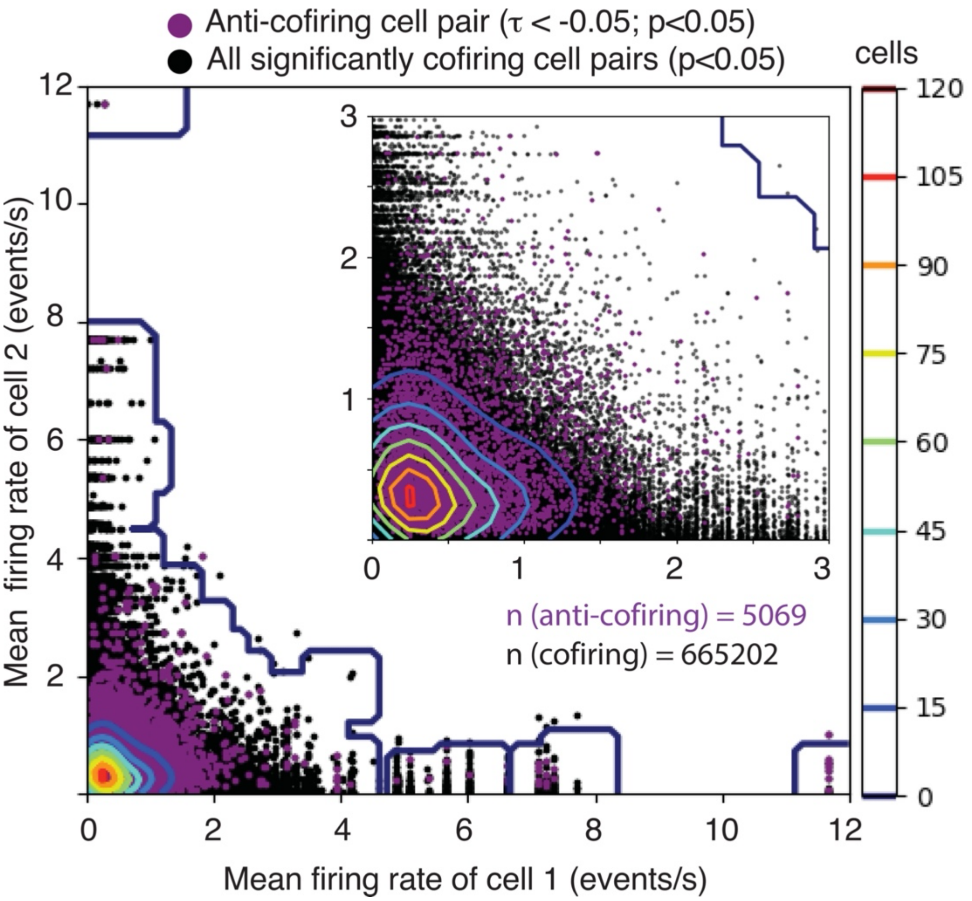


**Figure S9. Related to Figure 8. Pair-wise firing rate relationships of significantly anti-cofiring and cofiring cell pairs**. The overall activity of cells that constitute anti-cofiring cell pairs is typical of any cell pair, and does not mainly constitute cell pairs with only high and low activity rates. Density of cell pairs is indicated by the contour plots.

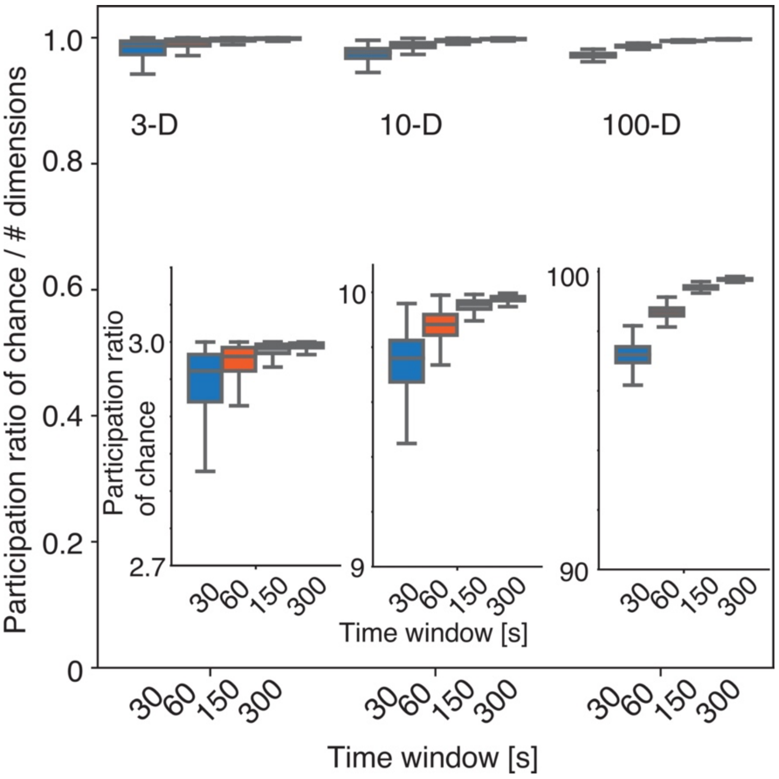


**Figure S10. Related to Figure 8. Dependence of participation ratio on sampling duration.** The time dependence of participation ration (PR) is substantially lost when PR is normalized by the dimensionality of the raw data. (inset) Participation ratio was computed for randomly selected 3-D, 10-D, and 100-D dimensional activity vectors of 300 sampled points, and across increasing time window durations of 30s, 60s, 150s and 300s.

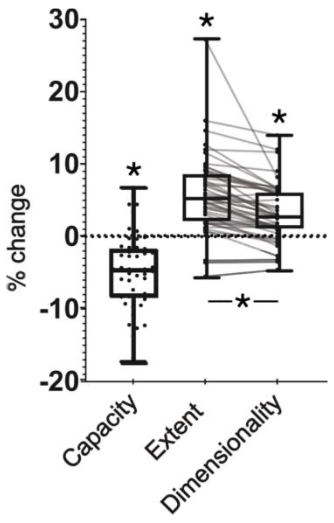


**Figure S11. Related to Figure 8. Geometric properties of high-dimensional neural manifolds from replica mean field theory of manifolds.** Relative change of the manifold geometric properties when removing the same proportion of anti-cofiring cells vs random cells. The capacity decreases (-5.13 ± 0.74; paired t test $t_{49}$ = 6.95, p = $10^{-8}$), meaning that the general overlap increases among manifolds. The relative increase in the extent metric (5.83 ± 0.76; paired t test $t_{49}$ = 7.69, p = $10^{-10}$) is also larger than the increase in dimensionality (3.27 ± 0.56; paired t test $t_{49}$ = 5.86, p = $10^{-7}$); a difference (paired t test $t_{49}$ = 6.14, p = $10^{-7}$) indicating that removal of the anti-cofiring cells reduced the distinctiveness more than it increases the dimensionality of the neuronal ensemble data.

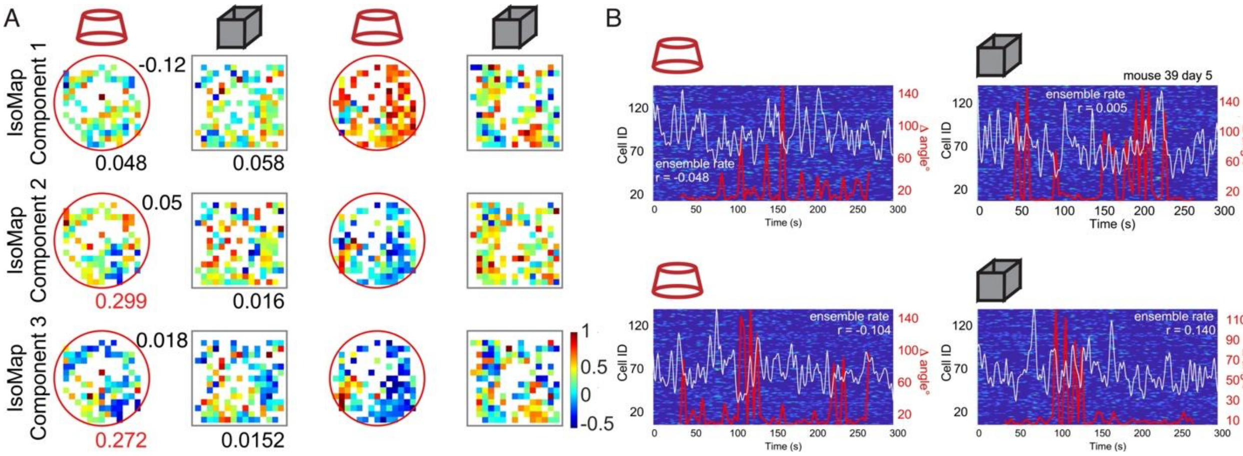


**Figure S12. Related to Figure 8. Spatial and temporal organization of manifold dynamics.** (A) Example average spatial distribution of IsoMap components. The time series of the first 3 IsoMap components (see Video S1) plotted as a session-averaged function of the mouse's position. The blue-to-red color-coded component map values are normalized to the maximum value observed during the day 5, 377-cell ensemble recording from mouse 39. Map correlations between select pairs of cylinder and box recordings are indicated below the first cylinder and box maps; the cylinder versus box correlation is given above between the two maps. Significant correlations are indicated in red. (B) No relationship was detected between the ensemble activity timeseries composed of the 134 most anti-coactive cells (white trace) and the 377-cell ensemble's planar manifold reorientations within the 3-D IsoMap subspace, measured as the timeseries of change in the 60-s planar orientation estimated each 4 seconds (red traces). The Pearson correlation between the white activity and red manifold timeseries is indicated. Single-unit ensemble activity timeseries is given in the blue-to-red background plot.

**Video S1. Related to Figure 7. CA1 ensemble activity vectors are constrained to environment-specific non-linear planar manifolds.** Video of 1-s activity vectors projected onto a 3-D subspace using the IsoMap non-linear dimensionality reduction algorithm, viewed from a rotating perspective. Individual points are projections of individual 1-s activity vectors and lines indicate average 3-D vector projections from each recording. Repetitions of the same environments on the same day are given distinct color shades.

**Video S2. Related to Figure 7. Dynamic jumbotron patterns as an example of an ensemble, non-linear manifold representation of an invariant message.** In this video of a jumbotron display the "Game Over" message is constant but which lights, the number of lights, where the lights turn on and off, which lights are on and off together and such, are all different, despite the message being the same. How CA1 activity represents a cylinder and a box lead us to consider that a manifold representation with no requirement for either single cell or ensemble stationarity is the appropriate conceptual framework for understanding how hippocampal activity represents spatial information. We analysed the 24-s video as a 180x320 (57600) ensemble time series across 720 frames (time steps), as described in Figures 6 and 7. The participation ratio that estimates the dimensionality is 600 (~1%). Projecting the 57600-D activity vectors into the 3-D non-linear subspace with the IsoMap algorithm (left), yields a reconstruction error increase of merely 5% relative to the reconstruction error using a 550-D IsoMap subspace, quantifying that most of the observed variance is captured by the 3 non-linear dimensions. A similar projection has been performed into the 3-D subspace using PCA (right). The two videos illustrate that the jumbotron's "Game Over" message occupies a very limited portion of the 3-D subspaces the axes of which are defined by particular jumbotron patterns of "Game Over."

**Video S3. Related to Figure 8. The CA1 ensemble activity vector timeseries evolves on transient, non-linear planar manifolds.** Video of 1-s activity vectors projected onto a 3-D subspace using the IsoMap non-linear dimensionality reduction algorithm. The dimensions (N1, N2, N3) that define the subspace were computed from a concatenated recording from a 475-cell recording in the cylinder and box from mouse 34 on day 5. The recording was 300-s in each environment. The animation illustrates the evolution of the 1-s activity vector positions in the 3-D IsoMap subspace for the cylinder (red) and box (gray). Once 60-s of data are accumulated the best fit plane is indicated and updated every 4-s. ΔAngle indicates the 4-s momentary change of the best fit plane's orientation in the subspace.
